# Predictive filtering of sensory response via orbitofrontal top-down input

**DOI:** 10.1101/2024.09.17.613562

**Authors:** Hiroaki Tsukano, Michellee M. Garcia, Pranathi R. Dandu, Hiroyuki K. Kato

## Abstract

Habituation is a crucial sensory filtering mechanism whose dysregulation can lead to a continuously intense world in disorders with sensory overload. While habituation is considered to require top-down predictive signaling to suppress irrelevant inputs, the exact brain loci storing the internal predictive model and the circuit mechanisms of sensory filtering remain unclear. We found that daily neural habituation in the primary auditory cortex (A1) was reversed by inactivation of the orbitofrontal cortex (OFC). Top-down projections from the ventrolateral OFC, but not other frontal areas, carried predictive signals that grew with daily sound experience and suppressed A1 via somatostatin-expressing inhibitory neurons. Thus, prediction signals from the OFC cancel out behaviorally irrelevant anticipated stimuli by generating their “negative images” in sensory cortices.

## Main Text

Throughout our lives, we are constantly exposed to a flood of sensory information from the external world. However, only a small fraction of this information reaches our conscious perception and triggers behavioral responses. This remarkable filtering, which prioritizes inputs with expected meaningful outcomes while ignoring others, arises from our brain’s ability to build internal models of the world (*1–3*). These models are continuously trained through our lifetime experiences and dictate predictive relationships between sensory objects in the world. The inability to build such predictive models, and thus to apply appropriate sensory filters, is a hallmark of mental disorders such as obsessive-compulsive disorder (*4*, *5*), schizophrenia (*6*), and most notably, autism spectrum disorders (ASD)(*7–10*).

One fundamental form of such sensory filtering is habituation, the brain’s ability to ignore familiar, inconsequential stimuli (*11–13*). Habituation operates through multitudes of mechanisms, allowing animals to adapt to the statistical structures of their sensory environments over a wide range of timescales. While previous research primarily focused on mechanisms for short-term plasticity lasting from milliseconds to seconds, known as “stimulus-specific adaptation” (*14–21*), habituation also extends to longer periods—days or even weeks (*22–26*)—as evidenced by the gradual reduction in our sensitivity to perfumes or traffic noise. This long-term habituation is particularly relevant for understanding the pathology of filtering deficits, as a lack of accumulated habituation to daily sensory objects throughout one’s life could result in a persistently intense sensory world (*27–32*).

Contrary to the widely held belief that habituation is the “simplest form of learning,” accumulating evidence has challenged this perceived simplicity. Classical internal model-free mechanisms, such as sensory receptor fatigue, homosynaptic depression, and the short-term dynamics of inhibitory neuron firing, fail to fully account for several key characteristics of habituation. These characteristics include its persistence over weeks, capacity to process complex stimuli involving multiple sensory channels, context-dependence (*33–35*), and disruption by anesthesia (*36*, *37*). These observations have prompted theories suggesting that habituation is associative and governed by top-down regulation informed by internal models of the external world (*11*, *29*, *36*, *38*, *39*).

Two competing hypotheses have been proposed to explain the top-down circuit mechanisms of long-term sensory habituation. The predictive negative image hypothesis posits that initially strong sensory responses are gradually canceled as the top-down predictive signal grows (Fig. 1A, left, and fig. S1A).

**Fig. 1.**
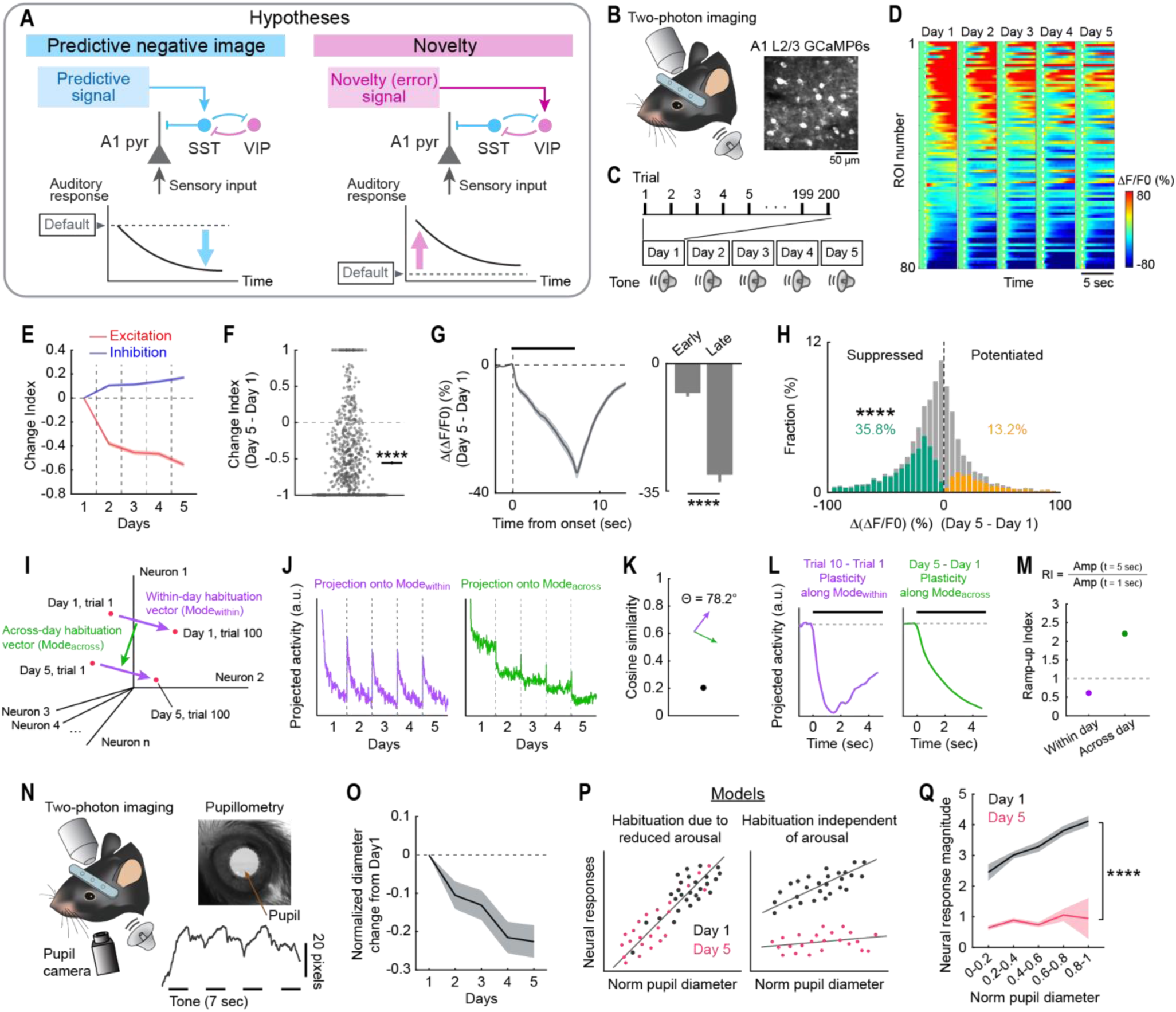
Inactivation of OFC reverses across-day habituation in A1. (**A**) Schematics illustrating two theories explaining sensory habituation by top-down predictive mechanisms. (**B**) Representative two-photon image of A1 L2/3 neurons expressing GCaMP6s. (**C**) Schematic illustrating the five-day auditory habituation paradigm. (**D**) Heatmaps of sound-evoked responses in neurons imaged across five days in a representative mouse. Neurons are sorted by their responses on Day 1. (**E**) Average (solid line) and SEM (shading) of the Change Index for excitatory (red) and inhibitory (blue) sound-evoked responses across days. *n* = 20 mice. (**F**) Change Index of excitatory responses in individual neurons on Day 5 compared to Day 1 across all mice. Black lines on the right represent mean ± SEM. *n* = 879 responsive neurons. \*\*\*\**P* = 1.6 × 10^-66^ (two-sided Wilcoxon signed-rank test). (**G**) Left: Change in sound-evoked response traces from Day 1 to Day 5 averaged across all significantly excited cells. Black bar: sound. Right: The amplitudes of the difference trace at 1 and 7 seconds after sound onset. \*\*\*\**P* = 1.4 × 10^-65^ (two-sided Wilcoxon signed-rank test). (**H**) Histograms of changes in response magnitudes in all neurons. Orange and green bars show neurons with significant increase and decrease. \*\*\*\**P* = 4.3 × 10^-138^ (Fisher’s exact test). (**I**) Schematic illustrating vectors representing within-day habituation (purple: Mode_within_) and across-day habituation (green: Mode_across_) within a high-dimensional space. Each dimension corresponds to the response magnitudes of individual neurons. (**J**) Projection of trial-to-trial sound-evoked A1 ensemble activity dynamics onto the Mode_within_ (left) and Mode_across_ (right) vectors. Data points represent individual trials (100 trials × 5 days). Ensemble activity patterns repeated fast daily plasticity along the Mode_within_ axis, while they show continuous slow plasticity along the Mode_across_ axis across five days. *n* = 20 mice, 2,398 cells imaged throughout the five days. (**K**) Cosine similarity between Mode_within_ and Mode_across_, indicating a nearly orthogonal relationship. (**L**) Left: Change in sound-evoked ensemble activity along Mode_within_ from Trial 1 to Trial 10, demonstrating prominent within-day, across-trial habituation around the tone onset. Right: Change in sound-evoked ensemble activity along Mode_across_ from Day 1 to Day 5. Across-day habituation slowly ramps up during the sustained tone. (**M**) Summary data showing the Ramp-up Index for across-day plasticity (Day 1 to Day 5) and within-day plasticity (Trial 1 to Trial 10). (**N**) Top left: Schematic illustrating pupil monitoring during two-photon calcium imaging. Top right: Representative image from a pupil camera. Bottom right: Representative pupil diameter dynamics during sound presentations. Black lines indicate tone timings. (**O**) Change in normalized pupil diameter from Day 1. *n* = 17 mice. (**P**) Two alternative models illustrating the dependence of neuronal sound responses on normalized pupil diameter. Left: A model in which neuronal habituation depends on the decrease in arousal level. Right: A model in which neuronal habituation is independent of arousal level. Individual dots represent trials. Black: Day 1 trials. Red: Day 5 trials. Lines indicate regression lines. (**Q**) Experimental data showing neuronal response magnitudes binned by normalized pupil diameter separately for Day 1 (black) and Day 5 (red). Day 5 neuronal responses are smaller than Day 1 responses regardless of normalized pupil diameter. Day 1 vs. Day 5: \*\*\*\**P* = 2.0 × 10^-30^ (unbalanced two-way ANOVA).

Supporting this hypothesis, somatostatin-expressing inhibitory neurons (SST cells) in sensory cortices increase their sensory responses over days of stimulus exposure (*40–42*), providing neuronal substrates for the generation of a ‘negative image’ of the stimulus. An alternative is the novelty (or prediction error) hypothesis, which assumes that sensory-evoked activity is weak by default and is amplified by a top-down novelty signal upon the animal’s first encounter with a stimulus (Fig. 1A, right)(*43–46*). This view proposes that the activation of vasoactive intestinal peptide-expressing inhibitory neurons (VIP cells) by the novelty signal disinhibits pyramidal neurons, and habituation occurs as the novelty signal wanes over time.

Despite their fundamentally distinct mechanisms, activity measurements in sensory cortices alone have failed to differentiate these hypotheses. Due to the reciprocal connections between SST and VIP cells, both models predict similar neuronal dynamics during habituation: an increase in SST cell activity and a decrease in VIP and pyramidal cell activity. To discriminate between these theories, we sought to identify the source of the top-down input to A1 during daily sensory habituation and determine the ‘default’ sensory responses by removing the top-down signal.

### A1 neurons show sound-specific habituation across days

We induced sensory habituation in awake, head-fixed mice through repeated exposure to tones over five days (5–9 seconds pure tones, 7–16 seconds inter-trial interval, 70 dB SPL, 200 trials/day) (Fig. 1, B and C). To track the activity of the same A1 layer 2/3 (L2/3) neural ensembles across days, we conducted chronic two-photon calcium imaging. We first located A1 by intrinsic signal imaging (*47*) and virally expressed GCaMP6s in the A1 of wild-type or VGAT-Cre × Ai9 mice. Three weeks after virus injection and optical window implantation, we recorded sound-evoked activities from 5,483 L2/3 neurons in 20 mice. We chose a fixed tone frequency for each mouse and imaged the corresponding tonotopic domain in A1, where we tracked the same neural ensembles throughout the experiment. Across days of sound exposure, we observed clear habituation in tone responses, characterized by reduced excitation and increased suppression in a sound-specific manner (Fig. 1, D to H, fig. S2, A to C, and fig. S3), consistent with previous studies (*40*, *41*). Given indistinguishable results between experiments imaging all L2/3 neurons and those imaging only VGAT^−^ pyramidal neurons, data from both experiments were combined and treated as A1 pyramidal cells (fig. S2, D to G).

We observed two timescales of habituation: within-day habituation, which repeated daily across trials, and across-day habituation, which built up over multiple days (Fig. 1, I and J). To better understand the relationship between these plasticity timescales, we projected high-dimensional population dynamics onto the axes defined by within-day and across-day changes. Notably, these two habituation mechanisms represented nearly orthogonal axes in the high-dimensional space (78.2 degree; Fig. 1K), indicating that across-day habituation is not simply the accumulated effect of within-day habituation. Within-day habituation markedly attenuated sound onset responses, whereas across-day habituation was greater during the sustained phase of tones (5 seconds after onset), implying two independent mechanisms for auditory habituation (Fig. 1, G, L, and M, and Supplementary Text). This difference in kinetics supports the role of slow, top-down predictive filtering in across-day habituation, while within-day habituation may be driven by faster, bottom-up mechanisms.

To assess the contribution of global neuromodulation to sensory habituation, we monitored pupil diameter during calcium imaging in a subset of experiments. Pupil diameter showed a gradual decrease across days, providing evidence of habituation in arousal responses (Fig. 1, N and O). However, two observations argued against the possibility that the A1 neuronal habituation stemmed from a general decrease in responsiveness due to lowered arousal. First, plots of trial-to-trial variability of neural activity against pupil diameter showed a distinct separation between Day 1 and Day 5 data regardless of pupil diameter, demonstrating that across-day habituation is independent of arousal levels (Fig. 1, P and Q). Second, independent calcium imaging using two different tone frequencies confirmed the sound specificity of A1 sensory habituation (fig. S3). These data suggest that sensory habituation in A1 is due to sound-specific plasticity rather than global brain state changes. Additionally, we previously found that across-day habituation under these conditions is not inherited from subcortical pathways, as the plasticity was evident in L2/3 but not in the input layer, L4 (*40*). Therefore, we focused on across-day habituation for the rest of our analyses to determine its cortical circuit mechanisms.

Both the predictive negative image hypothesis and the novelty hypothesis involve the active recruitment of inhibitory neurons during habituation. Since VIP cell activity had not been previously monitored in A1 during across-day habituation, we performed cell type-selective calcium imaging of three inhibitory neuron subclasses—SST cells, VIP cells, and parvalbumin-expressing neurons (PV cells)—to determine their dynamics over five days. We targeted GCaMP6s to each cell type by injecting a conditional GCaMP6s-expressing virus in Cre transgenic mice. Throughout the five-day sound exposure, L2/3 VIP and PV cells exhibited neural habituation similar to pyramidal cells (fig. S4).

Interestingly, SST cells showed clear heterogeneity (*48*); deep (>200 µm from the surface) SST neurons showed habituation similar to pyramidal cells, while superficial (<200 µm) neurons exhibited the opposite pattern (*40*, *41*). These findings suggest the involvement of the superficial SST cells in sensory habituation, consistent with the inhibitory dynamics proposed in both theories. However, they do not conclusively support one hypothesis over the other, prompting further investigation into the modulation source for A1 inhibitory neurons.

### OFC inactivation reverses sensory habituation in A1

We therefore sought to identify the source of top-down input regulating A1 neuronal activity (Fig. 2A). First, we performed anatomical tracing. After mapping auditory cortices with intrinsic signal imaging, we injected a retrograde tracer into A1 L2/3 (Fig. 2B). Consistent with previous studies, we found many presynaptic neurons in the OFC (*49–52*), secondary motor cortex (M2)(*53–55*), and anterior cingulate cortex (ACC)(*56*) (Fig. 2, C and D), with the ventrolateral and lateral OFC (hereafter, jointly called OFCvl) showing the densest inputs. The lack of overlap between the tracer and VGAT expression indicated that the top-down projections from OFCvl were glutamatergic (fig. S5, A to D). Cell type-selective retrograde tracing with rabies virus further confirmed that both SST and VIP cells receive extensive top-down inputs from OFCvl (Fig. 2, E to H, and fig. S6; see Discussion). In contrast, tracer signals in adjacent areas, such as the frontal pole (FRP), prelimbic area (PL), and infralimbic area (IL), were weaker in both AAV and rabies tracing, indicating highly specific wiring of auditory circuits in the frontal cortex (*57–59*).

**Fig. 2.**
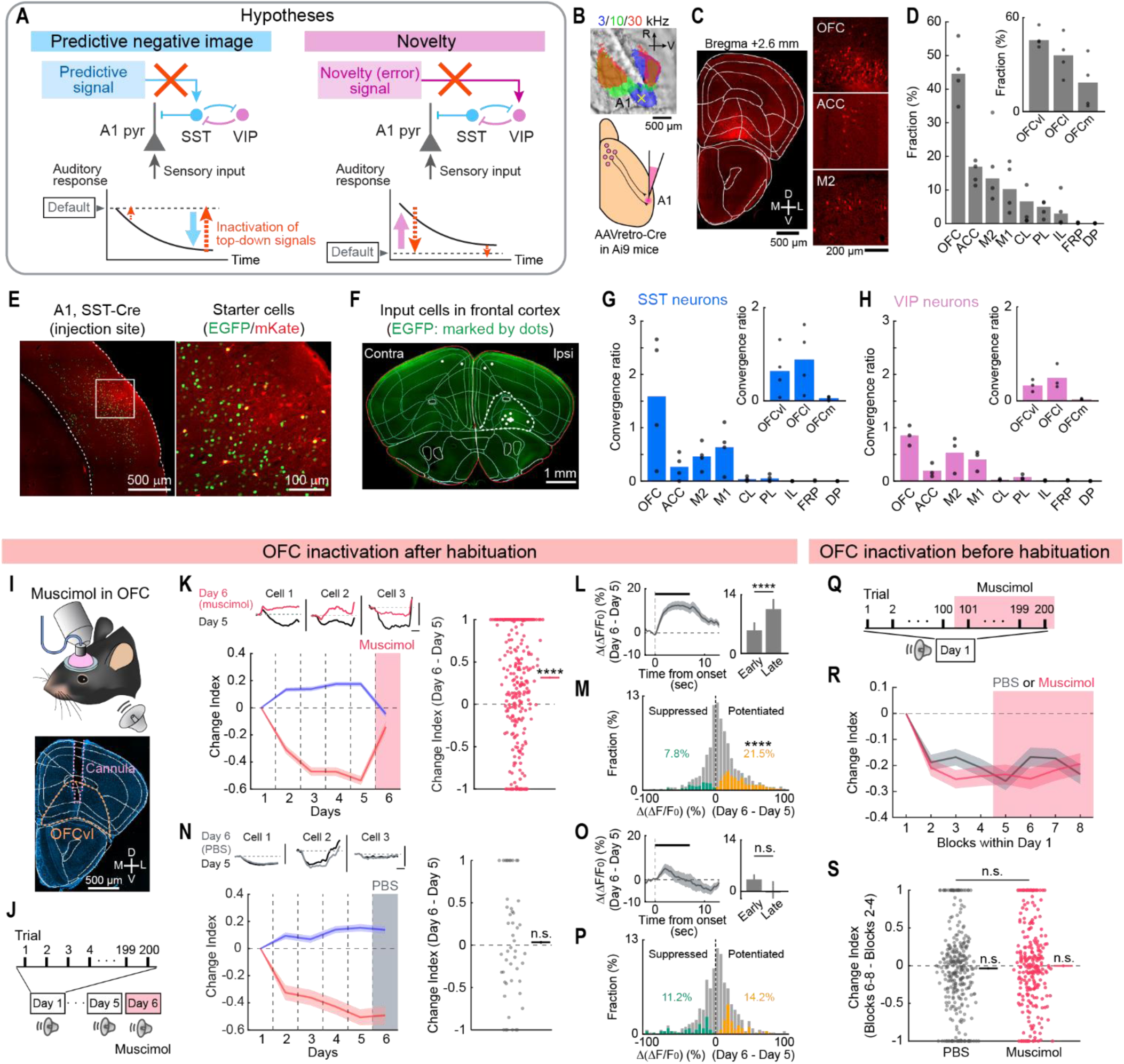
Inactivation of OFC reverses across-day habituation in A1. (**A**) Schematics illustrating two theories explaining how to distinguish top-down predictive mechanisms from novelty mechanisms. (**B**) Top: Intrinsic signal imaging of pure tone responses superimposed on the cortical surface imaged through the skull. Yellow cross: injection site. Bottom: Schematic illustrating retrograde tracing from A1 L2/3. (**C**) tdTomato-expressing input neurons onto A1 L2/3. Left: Coronal section of the frontal areas overlaid with area borders. Right: Magnified images of OFC, ACC, and M2 areas. (**D**) Bar plots summarizing the distribution of input cells in ipsilalteral frontal cortical areas. M1: primary motor cortex, CL: claustrum, DP: dorsal peduncular area. Inset: Further classification of OFC into ventrolateral (OFCvl), lateral (OFCl), and medial (OFCm) subdivisions. *n* = 4 mice. (**E-G**) Retrograde tracing from A1 L2/3 pyramidal neurons using SST-Cre mice. (**E**) Left: Coronal section of the injection site in A1. Dotted lines: cortical borders. Right: Magnified view of the area indicated by a square in the left image. (**F**) Coronal section of the frontal areas overlaid with area borders. White dots: presynaptic neurons. (**G**) Bar plots summarizing the distribution of input cells in ipsilateral frontal cortical areas. *n* = 4 mice. Inset: Further classification of OFC into ventrolateral (OFCvl), lateral (OFCl), and medial (OFCm) subdivisions. (**H**) Same as (G) but for VIP-Cre mice. (**I**) Top: Schematic illustrating muscimol infusion. Bottom: DAPI-counterstained coronal section showing the cannula implantation site in OFC. (**J**) Protocol for muscimol infusion after habituation. (**K**) Top left: Trial-averaged sound-evoked response traces from three cells. Bottom left: Change Index across days. Pink shade: Muscimol infusion. *n* = 7 mice, 1,392 neurons. Right: Change Index of excitatory responses in individual neurons on Day 6 compared to Day 5 across all mice. *n* = 296 significantly excited neurons. \*\*\*\**P* = 7.9 × 10^-16^ (two-sided Wilcoxon signed-rank test). (**L**) Left: Change in sound-evoked response traces from Day 5 to Day 6, averaged across all significantly excited cells. Right: The amplitudes of the difference trace at 1 and 7 seconds after sound onset. \*\*\*\**P* = 3.4 × 10^-7^. (**M**) Histograms of changes in response magnitudes in all neurons. \*\*\*\**P* = 5.7 × 10^-25^. (**N-P**) The same as (K-M), but for control PBS infusion experiments. (**N**) *n* = 3 mice, 482 total neurons, 49 significantly excited neurons. *P* = 0.52. (**O**) *P* = 0.23. (**P**) *P* = 0.19. (**Q**) Schematic illustrating muscimol infusion in naïve mice. (**R**) Change Index for excitatory responses across blocks of trials. Each data point represents a block of 25 trials. Red: muscimol (*n* = 4 mice). Dark gray: PBS control (*n* = 6 mice). (**S**) Change Index of excitatory responses in individual neurons in blocks 6–8 compared to blocks 2–4 across all mice. *n* = 301 and 242 significantly excited neurons for PBS and muscimol. PBS: *P* = 0.67 (two-sided Wilcoxon signed-rank test), muscimol: *P* = 0.23, PBS vs. muscimol: *P* = 0.49 (two-sided Wilcoxon rank-sum test).

We next asked whether the frontal top-down input causally contributes to A1 sensory habituation.

Two hypotheses predict distinct outcomes with the removal of top-down input (Fig. 2A, red dotted arrows). According to the predictive negative image model, removing the predictive signal should increase sensory responses following habituation but have minimal impact on naïve animals.

Conversely, the novelty model suggests that eliminating the novelty signal would diminish sensory responses in naïve animals but have little effect after habituation. To test these predictions, we targeted the OFCvl for pharmacological inactivation, given its strongest top-down connection to A1. After five days of tone habituation, we inactivated OFCvl on Day 6 by infusing muscimol through a pre-implanted cannula (2.5 mg/mL, 500 nL, 100 nL/min) (Fig. 2, I and J). Remarkably, OFCvl inactivation reversed the sensory habituation in A1 pyramidal cells by enhancing excitatory responses and reducing suppression responses, an effect not observed with the control PBS infusion (Fig. 2, K to P, and fig. S7) (Change Index from Day 5 to Day 6, muscimol: 0.31 ± 0.04, *P* = 7.9 × 10^-16^; control: 0.034 ± 0.094, *P* = 0.52). In a separate experiment, we evaluated the effects of OFCvl inactivation in naïve animals by infusing muscimol on Day 1 (Fig. 2Q). Unlike the post-habituation inactivation, OFCvl inactivation prior to habituation did not alter tone responses (Fig. 2, R and S) (Change Index before and after infusion, muscimol:-0.002 ± 0.017, *P* = 0.23; control:-0.037 ± 0.015, *P* = 0.67). The absence of enhanced responses by muscimol in naïve animals rules out the role of OFCvl in transmitting novelty signals or modulating general sensory processing in A1. Instead, our findings demonstrate the causal role of the OFC specifically in the expression of long-term sensory habituation, aligning with the predictive negative image hypothesis.

Given the involvement of specific inhibitory neuron subtypes in both hypotheses, we further assessed whether inhibitory neurons follow the predictions of these models. We repeated the same six-day OFCvl inactivation experiment while performing cell type-selective imaging of SST, VIP, and PV cells (Fig. 3A). Upon OFCvl inactivation in habituated mice, sound responses in SST cells were significantly reduced (Change Index from Day 5 to Day 6,-0.50 ± 0.06, *P* = 1.6 × 10^-9^), as expected from the predictive negative image hypothesis (Fig. 3, B to E). In contrast, both VIP and PV cells showed a marked enhancement in their sound responses, consistent with the release from SST cell-mediated inhibition (PV: 0.18 ± 0.06, *P* = 0.036; VIP: 0.55 ± 0.10, *P* = 0.0012) (Fig. 3, F to M, and fig. S7). These findings further support that the predictive signals from the OFC activate SST cells to suppress other A1 cell types after habituation.

**Fig. 3.**
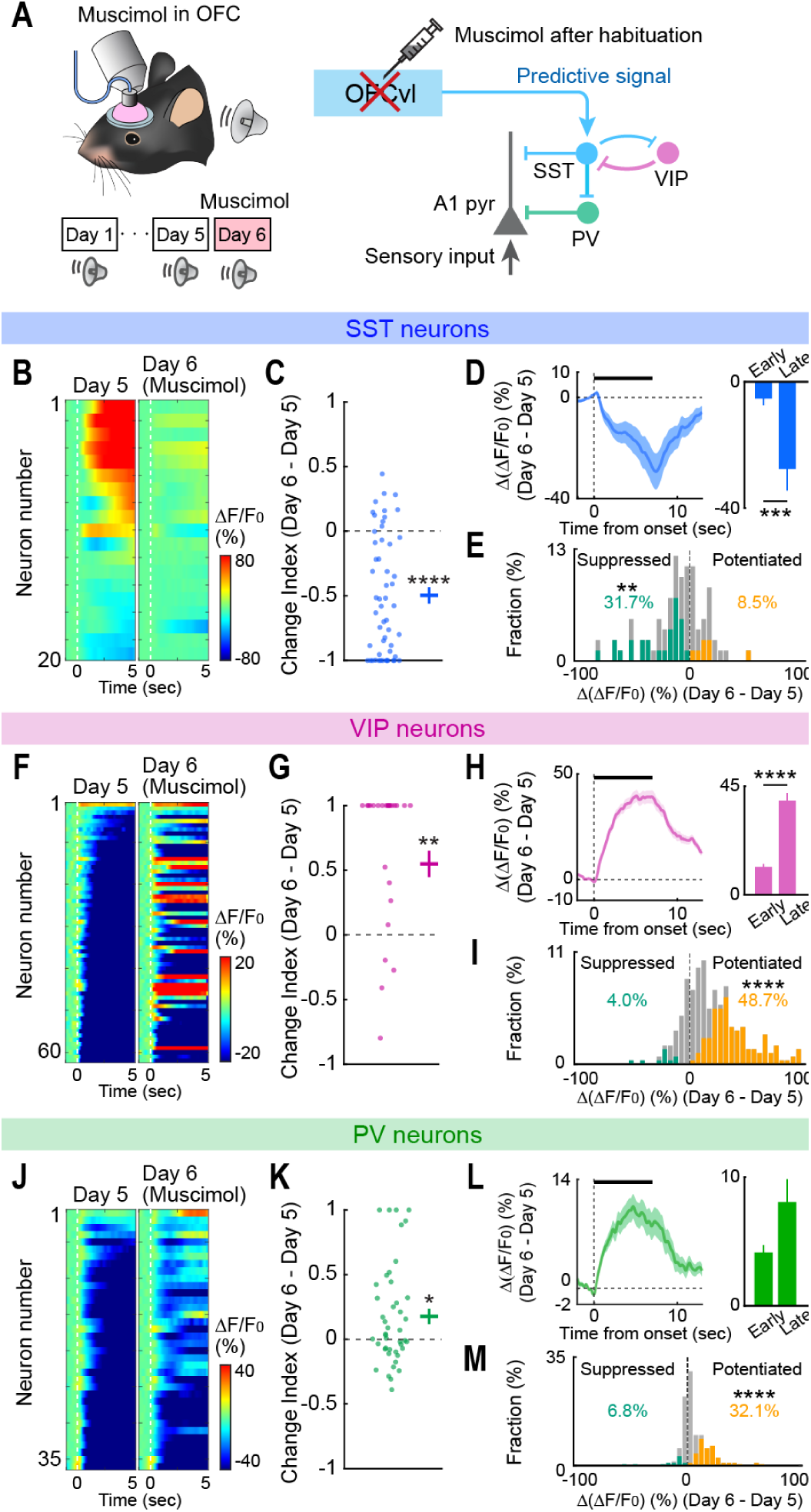
Post-habituation inactivation of OFC suppresses SST neuron activity in A1. (**A**) Left: Schematic illustrating muscimol infusion after habituation. Right: Circuit diagram of inhibitory neuron connectivity in A1. (**B**) Heatmaps of sound-evoked responses in SST neurons imaged across Days 5 and 6 in a representative mouse. Neurons are sorted by their responses on Day 5. (**C**) Change Index of excitatory responses in individual SST neurons on Day 6 compared to Day 5 across all mice. Lines on the right represent mean ± SEM. *n* = 4 mice, 54 excited cells. \*\*\*\**P* = 1.6 × 10^-9^ (two-sided Wilcoxon signed-rank test). (**D**) Left: Change in sound-evoked response traces from Day 5 to Day 6 averaged across all significantly excited SST cells. Solid line: average. Shading: SEM. Bar, sound. Right: The amplitudes of the difference trace at 1 and 7 seconds after sound onset. \*\*\**P* = 4.7×10^-4^ (two-sided Wilcoxon signed-rank test). (**E**) Histograms of changes in response magnitudes in all SST neurons. Orange and green bars show neurons with significant increase and decrease. \*\**P* = 0.0039 (Fisher’s exact test). (**F-I**) Same as (B-E) but for VIP neurons. (**G**) *n* = 4 mice, 26 excited cells. \*\**P* = 0.0012 (**H**) \*\*\*\**P* = 3.3 × 10^-26^. (**I**) \*\*\*\**P* = 3.3 × 10^-36^. (**J-M**) Same as (B-E) but for PV neurons. (**K**) *n* = 5 mice, 90 excited cells. \*\*\*\**P* = 7.9 × 10^-6^. (**L**) *P* = 0.30. (**M**) \*\*\*\**P* = 4.2 × 10^-5^.

Interestingly, while the effects of both sensory habituation (Fig. 1, G, L, and M) and OFCvl inactivation (Fig. 2, L, Fig. 3, D, H, and L, and fig. S7, E and F) were evident at sound onset (0–1 sec from the tone start), they were significantly more pronounced during the sustained sound stimulus. This slow kinetics of sensory habituation aligns with the recruitment of a long-distance top-down feedback loop and supports the role of prediction, as the sustained phase of the sound is more predictable than the onset.

### OFC top-down projection conveys predictive signals and suppresses A1

The OFC plays critical roles in various cognitive functions and has extensive connections with multiple brain regions, including the mediodorsal thalamus, striatum, amygdala, ACC, insular cortex, and sensory cortical areas (*60*). We next investigated whether the direct projection from OFCvl to A1 (fig. S5, E to H) conveys predictive signals during sensory habituation (Fig. 4A).

**Fig. 4.**
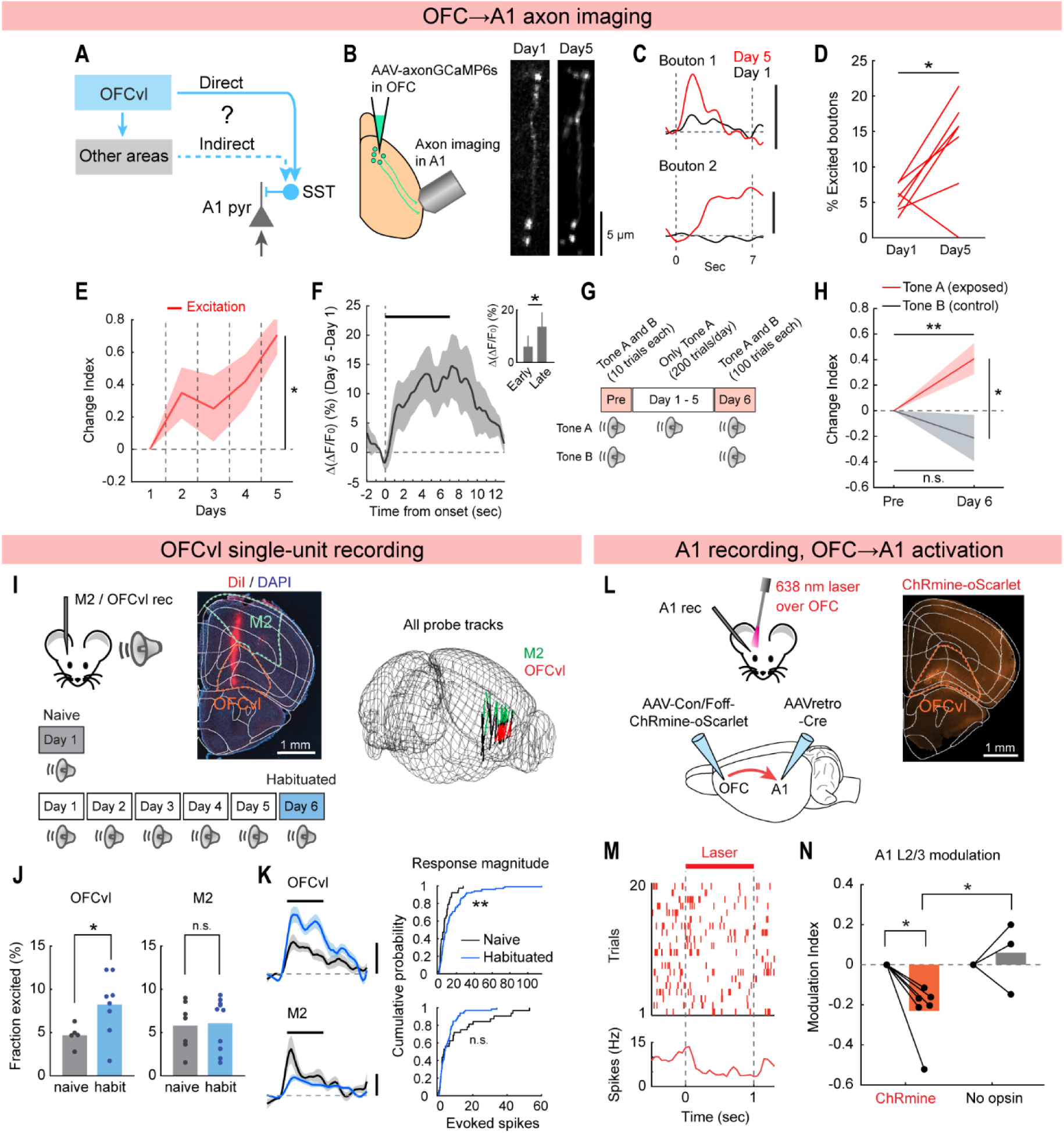
OFC top-down projection conveys predictive signals and suppresses A1. (**A**) Schematic illustrating direct and indirect pathways from OFC to A1. (**B**) Left: Schematic illustrating projection-specific calcium imaging of the OFC→A1 direct pathway. Right: Representative two-photon image of a GCaMP6s-expressing OFC axon in A1 on Day 1 and Day 5. (**C**) Trial-averaged sound-evoked response traces from two axon boutons. Black: Day 1. Red: Day 5. Scale bar: 50% ΔF/F. Dotted lines indicate sound onset and offset. (**D**) Fraction of boutons with significant excitatory responses on Day 1 and Day 5 across mice. *n* = 7 mice, 326 boutons. \**P* = 0.039 (two-sided Wilcoxon signed-rank test). (**E**) Average (solid line) and SEM (shading) of the Change Index for excitatory responses across days. \**P* = 0.016 (Day 1 vs. Day 5, two-sided Wilcoxon signed-rank test) (**F**) Change in sound-evoked response traces from Day 1 to Day 5 averaged across all significantly excited boutons. Black bar: sound. Inset shows the amplitudes of the difference trace at 1 and 7 seconds after sound onset. \**P* = 0.044 (two-sided Wilcoxon signed-rank test). (**G**) Schematic illustrating the sound-specific habituation paradigm using two sound frequencies. (**H**) Change Index of excitatory responses to Tone A (habituated) and Tone B (control) on Day 6 compared to the pre-habituation block. *n* = 7 mice, 18 and 13 significantly excited boutons for tones A and B. Tone A: \*\**P* = 0.0046 (two-sided Wilcoxon signed-rank test, Tone B: *P* = 0.20, Tone A vs. B: \**P* = 0.025 (two-sided Wilcoxon rank-sum test). (**I**) Left: Schematic illustrating single-unit recordings in M2 and OFC of naïve and habituated mice. Middle: Coronal section showing the Neuropixels probe track (indicated by DiI) targeting M2 and OFC. The section was counter-stained with DAPI and overlaid with area borders. Right: Summary of all penetrations aligned to the Allen Common Coordinate Framework. Probe tracks within M2 and OFC are shown in green and red, respectively. *n* = 20 penetrations in 17 mice. (**J**) Fraction of single units in OFCvl (left) and M2 (right) with significant excitatory responses in naïve and habituated mice. OFCvl naïve: *n* = 5 mice, habituated: *n* = 7 mice, \**P* = 0.045. M2 naïve: *n* = 7 mice, habituated: *n* = 10 mice, *P* = 0.83 (two-sided Wilcoxon rank-sum test). (**K**) Left: Sound-evoked response traces averaged across all significantly excited single units in OFCvl (top) and M2 (bottom) of naïve (black) and habituated (blue) mice. Right: Cumulative plots of sound-evoked spike counts in all significantly excited single units for OFCvl (top) and M2 (bottom). OFCvl, naïve: n = 38 units, habituated: n = 99 units, \*\**P* = 0.010. M2, naïve: *n* = 32 units, habituated: n = 66 units, *P* = 0.55. (**L**) Left: Schematic illustrating the intersectional strategy to express ChRmine selectively in A1-targeting OFC neurons. Right: Coronal section showing the expression of ChRmine-oScarlet. (**M**) Raster (top) and peristimulus time histogram (bottom) of a representative single unit in A1. (**N**) Modulation Index of A1 L2/3 neural activity by transcranial photostimulation of OFC with a red laser. ChRmine: *n* = 6 mice, *P* = 0.012 (two-sided Wilcoxon signed-rank test). No opsin control: *n* = 3 mice, *P* = 0.67. ChRmine vs. no opsin: *P* = 0.034 (two-sided Wilcoxon rank-sum test).

We visualized sound-evoked activity in the OFCvl→A1 direct pathway by virally expressing axonGCaMP6s in OFCvl neurons. We performed two-photon calcium imaging of their axon terminals in L1 and superficial L2/3 of A1 to track the activity from the same axon boutons over a five-day habituation period (Fig. 4B). On Day 1, significant tone-evoked excitation was observed in 5.5% of axon boutons, consistent with the significant but sparse sound responses previously reported in OFC of naïve mice (*49*, *61*) (Fig. 4, C and D). The sound responses are likely conveyed through the anatomical projections from the auditory cortex to the OFCvl (fig. S5, I to O). However, following habituation, the proportion of excited boutons more than doubled to 13.2%. In axon boutons identified across days, we observed a gradual increase in response magnitudes (Fig. 4, C and E). This increase in OFCvl→A1 axonal activity was more pronounced during the sustained phase of tones (Fig. 4F), consistent with the slow kinetics of habituation in A1 neurons (Fig. 1G, and Fig. 3, D, H, and L). Moreover, this plasticity was specific to the habituated tone frequency, ruling out a general increase in OFCvl responsiveness (Fig. 4, G and H). These results support the hypothesis that the OFC gradually forms an internal model of repeated stimuli, and its direct projection to A1 carries predictive signals that grow over days to recruit SST inhibitory neurons.

Next, we examined whether the increase in the representation of habituated sounds is specific to the OFCvl. Previous research found predictive filtering of A1 responses to self-generated sounds by M2→A1 top-down input (*53–55*). Thus, predictive filtering of sensory responses may be a general function shared equally across frontal cortical areas. Alternatively, distinct frontal regions might apply unique filters on A1 activity, tailored to different types of predictions: OFC for predictions based on a history of experiences, and M2 for predictions related to motor corollary discharges.

To investigate these possibilities, we directly compared tone-evoked activities between OFCvl and M2 neurons within the same animals. We performed single-unit recordings using a Neuropixels 1.0 probe that penetrated both regions (Fig. 4I). When we compared tone responses between naïve and habituated mice, we found a 1.7-fold larger fraction of significantly excited OFCvl single units in habituated mice (habituated: 8.2%; naïve: 4.7%, *P* = 0.045), while M2 single-units showed no difference (habituated: 6.1%; naïve: 5.8%, *P* = 0.83) (Fig. 4J). Sound-evoked response magnitudes averaged across all excited single-units showed a significant increase after habituation in OFCvl (104% increase; *P* = 0.010; Fig. 4K). In contrast, M2 units showed a tendency for reduced activity, likely reflecting a weaker input from the habituated auditory cortex (47% reduction; *P* = 0.55). Recordings in other frontal regions also showed no signs of increased tone responses (fig. S8, A to F). This OFCvl-specific plasticity is consistent with the OFCvl→A1 axonal imaging data and supports a division of labor between OFC and M2 in the predictive filtering of A1 activity across different contexts.

To determine whether activation of the OFCvl→A1 pathway is sufficient to suppress A1 neuronal activity, we next performed pathway-selective optogenetic activation. We restricted the expression of red-shifted opsin ChRmine in A1-projecting OFCvl neurons by an intersectional viral approach (Fig. 4L). Transcranial illumination of the frontal cortex with a red laser successfully evoked action potentials in ChRmine-expressing OFCvl neurons (fig. S8, G and H)(*62*, *63*). A1 recording during activation of OFCvl→A1 pathway revealed significant suppression of A1 activity, an effect absent in control mice without opsin expression (Modulation Index, ChRmine: −0.23 ± 0.06, *P* = 0.012; Control: 0.05 ± 0.07, *P* = 0.668; ChRmine vs. Control: *P* = 0.034; Fig. 4, M and N). Together, these results indicate that the OFCvl stores the information of repeated sensory stimuli and directly transmits predictive signals to A1, triggering neural habituation in A1.

### Plasticity in SST cells amplifies sensory habituation

Our results have demonstrated neural plasticity in the OFC, leading to the predictive filtering of A1 sensory responses. However, the OFC is considered to support flexible learning not only through its own plasticity (*64–67*) but also by driving plasticity in other brain regions (*68–71*). A previous study found that pairing OFC→A1 axon terminal activation with tone presentations results in plastic changes in A1 that suppress its responses to the paired frequency (*49*). Therefore, we wondered whether the enhanced OFCvl→A1 input during habituation triggers synaptic plasticity in A1 to further amplify sensory habituation.

We first explored the involvement of overall A1 synaptic plasticity in sensory habituation through the area-selective knockout of N-methyl-D-aspartate (NMDA) receptors. We co-injected AAV9-GCaMP6s and AAV8-mCherry-Cre into the A1 of floxed GluN1 mice (Grin1^flox/flox^) (Fig. 5A). Previous studies using the same mouse showed that Cre-dependent knockout of NMDA receptors became evident within two weeks post-transfection and grew over subsequent weeks (*72–74*). Five weeks post-injection, tone response magnitudes of A1 pyramidal cells were smaller in GluN1 knockout mice compared to control mice (See Methods) (fig. S9, A to D), consistent with the loss of NMDA receptors. Comparing five-day sensory habituation of GluN1 knockout mice to controls revealed a significant reduction in across-day sensory habituation (Fig. 5B), suggesting that local synaptic plasticity in A1 amplifies long-term sensory habituation.

**Fig. 5.**
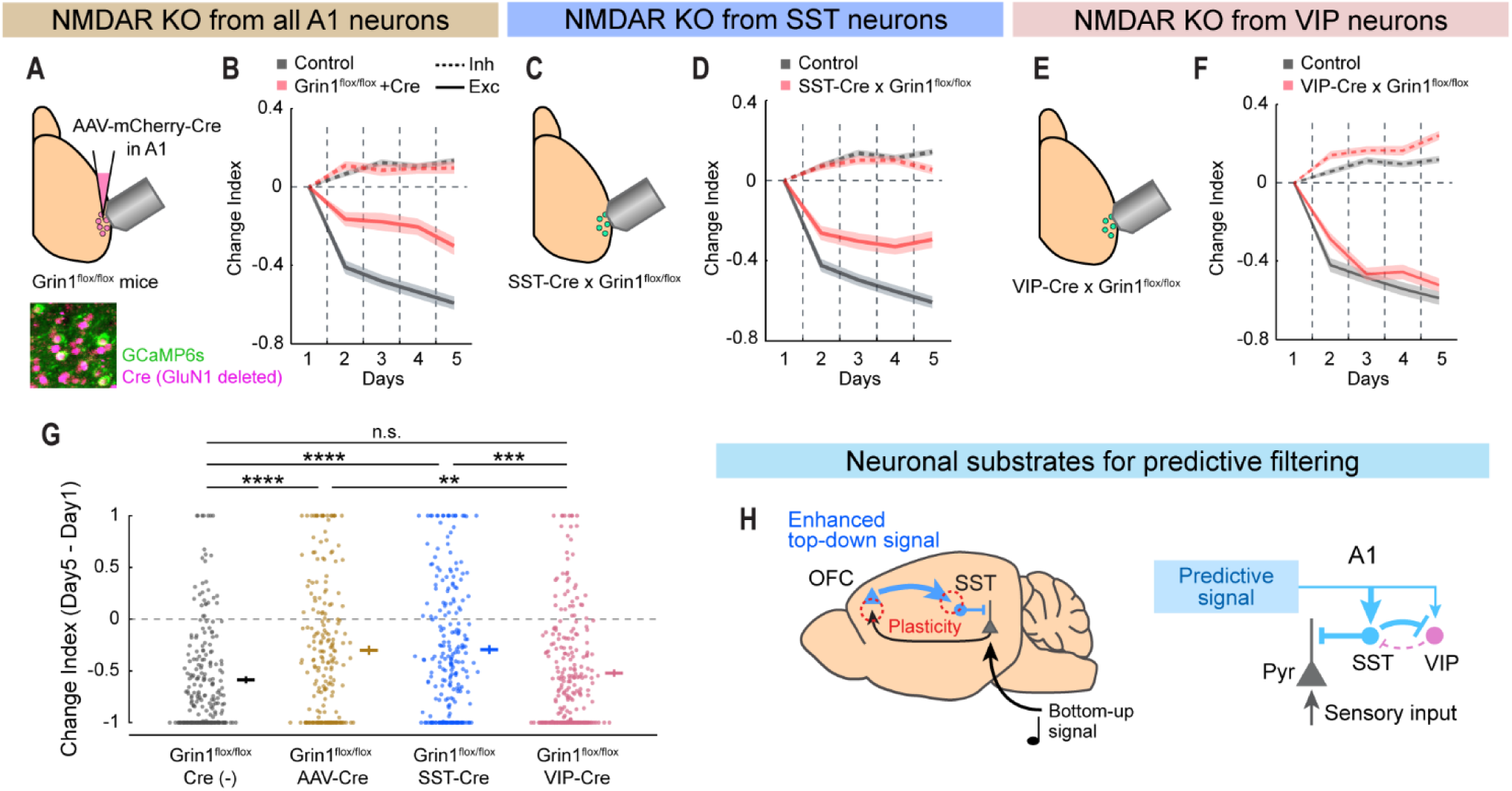
NMDA receptors in A1 and SST cells are required for habituation. (**A**) Top: Schematic illustrating the viral strategy to knock out NMDA receptors in A1 neurons. Bottom: Representative two-photon image of A1 L2/3 neurons from a Grin1^flox/flox^ mouse expressing GCaMP6s (green) and Cre (red). (**B**) Change Index for excitatory (solid line) and inhibitory (dotted line) sound-evoked responses across days in Grin1^flox/flox^ (red) and control (black) mice. Grin1^flox/flox^: *n* = 4 mice, 905 cells. Control: *n* = 5 mice, 1,708 cells. (**C-D**) Same as (a-b) but for knockout of NMDA receptors selectively in SST neurons. (**E**) SST-Cre × Grin1^flox/flox^: *n* = 5 mice, 1,364 cells. Control: *n* = 6 mice, 1,740 cells. (**E-F**) Same as (a-b) but for knockout of NMDA receptors selectively in VIP neurons. (**F**) VIP-Cre × Grin1^flox/flox^: *n* = 4 mice, 996 cells. Control: *n* = 5 mice, 1,494 cells. (**G**) Summary scatter plots showing the Change Index of excitatory responses in individual neurons on Day 5 compared to Day 1. Grin1^flox/flox^, Cre^−^ control (Ctrl): *n* = 226 excited cells. Grin1^flox/flox^ + AAV-Cre (KO_A1_): *n* = 210. SST-Cre × Grin1^flox/flox^ (KO_SST_): *n* = 231. VIP-Cre × Grin1^flox/flox^ (KO_VIP_): *n* = 254. Ctrl vs. KO_A1_: \*\*\*\**P* = 6.9 × 10^-5^. Ctrl vs. KO_SST_: \*\*\*\**P* = 3.2 × 10^-7^. Ctrl vs. KO_VIP_: *P* = 1.0. KO_A1_ vs. KO_SST_: *P* = 1.0. KO_A1_ vs. KO_VIP_: \*\**P* = 0.0034. KO_SST_ vs. KO_VIP_: \*\*\**P* = 3.2 × 10^-4^. (two-sided Wilcoxon rank-sum test with Bonferroni correction). (**H**) Summary diagrams of sensory habituation pathways.

Finally, we focused on synaptic plasticity specifically in SST cells. Given that GluN1 is also expressed in SST cells and regulates their plasticity (*75*, *76*), we generated Grin1^flox/flox^ mice carrying SST-Cre to delete GluN1 selectively in SST cells. A1 pyramidal cells in these mice showed normal tone response properties in a naïve condition (fig. S9, H to K). However, quantification of tone responses across days revealed a significantly diminished habituation in SST cell GluN1 knockout mice compared to the control group (Fig. 5, C and D). In contrast, selective deletion of GluN1 in VIP cells did not affect sensory habituation (Fig. 5, E to G). Thus, while GluN1 deletion in SST cells does not alter basal sound processing in A1, it attenuates sensory habituation through decreased synaptic plasticity. Collectively, our experiments demonstrate that experience-dependent plasticity in both OFCvl circuits and A1 SST cells leads to sensory habituation via the recruitment of SST inhibitory circuits by top-down predictive signals.

## Discussion

Theories based on human psychophysics have suggested the role of internal predictive models in sensory habituation (fig. S1). Our findings identify the neural substrates for these predictive models and the negative images of sensory stimuli they generate. In naïve animals, bottom-up sensory signals evoke strong responses in the A1. However, with repeated exposure to the same sound over days, a predictive model forms in the OFC. The OFC then sends top-down predictive signals to SST cells in the A1, which in turn suppress the activity of pyramidal, VIP, and PV cells. This finding may appear at odds with the focus of previous studies on the top-down regulation of VIP cells and the recruitment of the VIP→SST disinhibitory circuit (*48*, *77*, *78*). However, quantification of cell type-specific retrograde tracing from sensory cortical areas has revealed denser top-down connectivity from frontal cortical areas to SST than VIP cells (*52*) (fig. S6). Furthermore, SST and VIP cells are known to form a mutual inhibitory loop (*79–81*), with SST→VIP connections exhibiting one of the highest connection probabilities, largest IPSP amplitudes, and strongest facilitation within cortical circuits (*82*). Therefore, it is not unexpected that top-down input from OFCvl, which connects to both SST and VIP cells, could engage the often overlooked SST→VIP inhibitory connection to suppress the A1 circuit. Synaptic plasticity in SST neurons may further shift the balance of SST-VIP mutual inhibition toward SST dominance to facilitate habituation (Fig. 5H). Our model successfully explains recent studies in V1 that identified a VIP-dominant disinhibitory circuit during animals’ first encounter with novel stimuli (*43*, *44*, *83*). We do not exclude the roles of novelty or prediction error signals in other contexts (Supplementary Discussion).

However, our data indicate that daily sensory habituation is mediated by an increase in predictive signals, rather than a decrease in novelty (prediction error) signals. Thus, plasticity in the OFC, through the accumulation of experiences over extended periods (*64–67*), creates and updates the internal predictive model of the sensory environment, which is then used to filter neural activity in sensory cortices.

The OFC is considered to support flexible behaviors by encoding the values of predicted outcomes (*84–86*), assigning prediction error credits to corresponding actions (*87*), and tracking associative relationships between stimuli (*1*, *2*, *69*, *88*). However, the extent to which the OFC’s top-down projections influence sensory processing remains debated (*49*, *50*, *68*, *89*). Our finding of the OFC’s role in the predictive filtering of sensory inputs aligns with the concept that the OFC constructs and uses internal associative models, or “cognitive maps,” which dictate predictive relationships between the sensory task space and possible outcomes (*1*, *2*). Furthermore, our results suggest a more generalized function of the OFC beyond merely predicting explicitly rewarding or punishing action outcomes, extending to the attenuation of salience in neutral stimuli during passive experiences. This broader role is reminiscent of the OFC’s involvement in “devaluation,” where initially favorable stimuli lose their value with satiety or new associations with unfavorable outcomes (*90–92*). It is plausible that sensory inputs inherently carry behavioral values, as they may signal the approach of rewards or threats crucial for an animal’s survival. By forming internal associative models, the OFC “devalues” not only explicitly rewarding stimuli but also neutral sensory inputs without meaningful outcomes, thereby inducing habituation.

It is worth pointing out the similarity between our findings and the predictive filtering of self-generated sounds, which has been proposed to involve the top-down pathway from the M2 to the A1 after days of movement-sound association (*53*, *54*). These studies suggested the recruitment of PV inhibitory neurons, which broadly scale down the A1 activity during movements. Although exact circuits are different, these findings collectively support a general principle: frontal cortical areas send top-down predictive signals based on days of experience—M2 for movement-related predictions and the OFC for familiarity-based predictions, respectively—to filter predictable sensory inputs. In support of this division of labor, we observed a selective enhancement of sound-evoked activity in the OFC, but not in M2, following sensory habituation. Aside from the predictive coding, previous studies also suggested attentional filtering of A1 activity by the prefrontal cortex (*93–96*). Understanding how individual frontal areas are recruited in specific contexts to apply these appropriate filters would be an important area for future research.

Impaired habituation to repeated stimuli and resultant sensory hypersensitivity are commonly observed in mental disorders such as obsessive-compulsive disorders (*4*, *5*) and ASD (*7–10*). Although theories have proposed that a lack of predictive filtering may be central to the symptoms associated with ASD (*29–31*), the neural substrates linking altered neural wiring with reduced sensory filtering have remained unclear. Our discovery that top-down projections from the OFC to A1 are essential for auditory habituation provides crucial insights into the circuit mechanisms underlying reduced habituation in individuals with ASD. It is believed that the neocortical network in ASD displays long-range hypoconnectivity, as indicated by less cohesive activity across brain regions and reduced white matter volume (*97–100*). This reduced long-range communication between frontal areas and sensory cortices could therefore be a culprit for the insufficient habituation in these disorders. This general framework of top-down predictive filtering highlights the importance of targeting long-range frontal projections in developing interventions to alleviate sensory pathophysiology.

## Acknowledgments

We thank Paul Manis, Jeffry Isaacson, Daniel Polley, and the members of the Kato Lab for their advice throughout the project and comments on the manuscript. We thank Gonçalo Lopes for his support in setting up Bonsai for pupil monitoring.

## Author contributions

Conceptualization: HT, HKK

Methodology: HT, MMG, HKK

Investigation: HT, MMG, PRD, HKK

Formal analysis: HT, HKK

Visualization: HT, HKK

Funding acquisition: HT, HKK

Project administration: HKK

Supervision: HKK

Writing – original draft: HT, HKK

Writing – review & editing: HT, MMG, PRD, HKK

## Data and materials availability

All data generated in this study will be deposited in the DANDI Archive. Source data for all figures will be provided in this paper as a supplementary data file attached to this paper. Custom Matlab codes used in this study will be made available from the corresponding author upon request.

