## Supplementary Information for "Predictive filtering of sensory response via orbitofrontal top-down input"

**Materials and Methods:**

<u>Animals</u>

Mice were at least 6 weeks old at the time of experiments. The strains used include: C57BL/6J (JAX000664), B6N-Cdh23$^{\text{tm2.1Kjn}}$/Kjn (JAX 018399), Slc32a1$^{\text{tm2(cre)Lowl}}$/J (VGAT-Cre; JAX 028862), Pvalb$^{\text{tm1(cre)Arbr}}$/J (PV-Cre; JAX 017320), Sst$^{\text{tm2.1(cre)Zjh}}$/J (Sst-Cre; JAX 013044), Vip$^{\text{tm1(cre)Zjh}}$/J (VIP-Cre; JAX 010908), Gt(ROSA)$^{\text{26Sortm9(CAG-tdTomato)Hze}}$/J (Ai9; JAX 007909), and B6.129S4-Grin1$^{\text{tm2Stl}}$/J (Grin1$^{\text{flox/flox}}$; JAX 005246). Both female and male mice were used. The animals were housed at 21°C and 40% humidity, with ad libitum access to food pellets and water. They were kept under a reverse light cycle (12–12 h), and all experiments were conducted during their dark cycle. All experimental procedures were approved and conducted in accordance with the Institutional Animal Care and Use Committee at the University of North Carolina at Chapel Hill and the guidelines of the National Institutes of Health.

<u>Auditory stimuli</u>

Auditory stimuli were calculated in Matlab (MathWorks) at a sample rate of 192 kHz and delivered via a free-field electrostatic speaker (ES1; Tucker-Davis Technologies). The speakers were calibrated across a range of 2–64 kHz to give a flat response (±1 dB). Stimuli were delivered to the ear contralateral to the imaging or recording site. The delivery of auditory stimuli was controlled by Bpod (Sanworks), running on Matlab.

For areal mapping with intrinsic signal imaging, 3, 10, and 30 kHz pure tones (75 dB SPL, 1-second duration) were presented at 30-second intervals for 5–20 trials. The tonal receptive fields of individual neurons were determined by presenting 1-second pure tone pips covering 17 frequencies (4–64 kHz, log-spaced) at 30, 50, and 70 dB SPL. Tone pips were presented in a randomized order at 5-second intervals, with each stimulus repeated across five trials. For auditory habituation experiments, a single frequency was chosen for each mouse (see Two-photon calcium imaging section). Prolonged pure tones (70 dB SPL,

5, 7, or 9-second duration) were presented at 7–16-second intervals, 100 or 200 trials per day. For linear probe recordings in A1, the cortical layers were identified through current source density analysis using click sounds, which were generated as 0.1-ms monopolar rectangular pulses and presented across 200 trials at 0.5-second intervals. All pure tone stimuli had a 5-ms linear rise and fall at their onset and offset.

<u>Intrinsic signal imaging</u>

Intrinsic signal imaging was conducted to locate the auditory cortical areas, following a previously described protocol (*47*). Signals were acquired using a custom tandem lens macroscope comprising Nikkor 35 mm 1:1.4 and 135 mm 1:2.8 lenses, attached to a 12-bit CCD camera (DS-1A-01M30, Dalsa). Mice were anesthetized with isoflurane (0.8–2%) vaporized in oxygen (1 L/min) and maintained on a heating pad at 36°C. The muscle overlying the right auditory cortex was removed, and a custom stainless steel head bar was secured to the skull with dental cement. For initial mapping prior to craniotomies, the brain surface was imaged through a skull kept transparent with phosphate-buffered saline (PBS). For re-mapping before two-photon calcium imaging, the brain surface was imaged through an implanted glass window. Mice received a subcutaneous injection of chlorprothixene (1.5 mg/kg) prior to imaging. Images of surface vasculature were captured under green LED illumination (530 nm), and intrinsic signals were recorded at 16 Hz using red illumination (625 nm). Images of reflectance were acquired at a resolution of 717 × 717 pixels (2.3 × 2.3 mm). Images collected during the response period (0.5–2 seconds from sound onset) were averaged across trials and normalized to the average image during the baseline period. The images were Gaussian filtered and thresholded for visualization. Individual auditory areas, including A1, AAF, VAF, and A2, were identified based on their characteristic tonotopic patterns. Mice with distorted maps or missing responses were excluded before experiments.

<u>Two-photon calcium imaging</u>

Following initial transcranial mapping of auditory cortical areas using intrinsic signal imaging, a craniotomy (2 × 3 mm) was performed over the right auditory cortex, leaving the dura intact. Except for axon terminal imaging, viruses were injected using sharpened glass pipettes (15–20 μm outer diameter) at 7–10 locations around A1 (180–250 μm deep from the pial surface, 30–40 nL/site at 10 nL/min). For pyramidal cell imaging, AAV9-syn-GCaMP6s was injected in C57BL/6J or VGAT-Cre×Ai9 mice. For interneuron subtype-selective imaging, AAV9-syn-Flex-GCaMP6s was injected in either PV-Cre, SST-Cre, or VIP-Cre mice. For imaging with A1-selective NMDA receptor knockout, AAV9-syn-GCaMP6s and AAV8-mCherry-Cre were co-injected in Grin1$^{flox/flox}$ mice. For cell type-selective knockout of NMDA receptors, AAV-9-syn-GCaMP6s was injected in SST-Cre×Grin1$^{flox/flox}$ or VIP-Cre×Grin1$^{flox/flox}$ mice. For imaging OFCvl axon terminals in A1, AAV1-syn-axonGCaMP6s was stereotaxically targeted to the right OFCvl at two coordinates: (Anterior, Lateral, Ventral) = (2.4, 0.85, 2.5) and (2.7, 0.85, 2.5) from bregma in millimeters (300 nL/site at 20 nL/min). A glass window was then placed over the craniotomy and secured with dental cement. Dexamethasone (2 mg/kg) was injected prior to the craniotomy. Enrofloxacin (10 mg/kg) and meloxicam (5 mg/kg) were administered before the mice were returned to their home cage.

Two-photon calcium imaging was conducted 2–7 weeks after chronic window implantation to ensure an appropriate level of GCaMP6s expression. A second intrinsic signal imaging through the chronic window was conducted 1–3 days before two-photon calcium imaging to confirm the integrity of the tonotopic map. During calcium imaging, awake mice were head-fixed under a two-photon microscope in a custom-built sound-attenuation chamber. One to two fields of view (FOVs) in A1 were selected based on intrinsic signal maps. Before starting habituation experiments, tonal receptive fields of individual neurons were determined by measuring responses to pure tone pips. After constructing the best frequency maps, one FOV was selected in the low-to-middle-frequency domain of A1 to avoid the influence of high-frequency hearing loss. Sound frequencies for passive exposure (4–22 kHz) were chosen in each mouse as the frequency that evoked excitation in the largest fraction of cells in the imaged FOV. For sound-specific habituation experiments, two frequencies with a one-octave separation flanking the best frequency of the FOV were selected to ensure comparable response magnitudes in naïve conditions. During habituation, mice underwent either a single day or five consecutive days of imaging, with a pure tone presented 100 or 200 trials per day. GCaMP6s and tdTomato were excited at 925 nm, and mCherry was excited at 1070 nm (InSight DS+, Newport). Images (512 × 512 pixels covering 620 × 620 μm for imaging soma and 124 × 124 μm for imaging axon terminals) were acquired at 30 Hz using a commercial microscope (MOM scope, Sutter) equipped with a ×16 objective (Nikon) and running Scanimage software (Vidrio). Imaging was conducted in L2/3 (120–280 μm below the surface) for soma and in L1 and shallow L2 (60–130 μm below the surface) for axon terminals.

Muscimol infusion

For the pharmacological inactivation of the OFC, a cannula was implanted during the same surgery as the craniotomy. A guide cannula (outer diameter 0.41 mm, inner diameter 0.25 mm; RWD) was stereotaxically inserted at coordinates (A, L, V) = (2.7, 0.85, 2.7) in the right hemisphere. The guide cannula was secured with dental cement and sealed with a dummy cannula. On the day of muscimol infusion, the dummy cannula was replaced with an internal cannula (outer diameter 0.21 mm, inner diameter 0.11 mm; RWD) containing either a muscimol solution (2.5–5 mg/mL in PBS) or PBS (for control). A total volume of 500 nL of the solution was infused at 100 nL/min using a micropump (UMP3 and Micro4, WPI). Infusions were carried out before starting Day 6 of sound exposure (habituated mice) or after 100 trials of sound exposure on Day 1 (naïve mice).

Pupillometry

The eye contralateral to the craniotomy was imaged using an infrared camera (Genie Nano M640-NIR, Teledyne Dalsa, or Firefly FFY-U3-16S2M-S, FLIR). The two-photon excitation laser transmitted through the eye was used to visualize the pupil. A violet LED (395 nm) provided low-intensity illumination, maintaining the pupil at approximately mid-range in diameter. The intensity and position of the LED were kept consistent across chronic imaging sessions to allow for cross-day comparisons of pupil diameters. Pupil movies were captured and synchronized with sound presentation trials using Bonsai software (https://bonsai-rx.org/). Images of 360 × 480 pixels were collected at 30 Hz. The pupil diameter was measured using a custom Matlab program. First, an intensity threshold was set for each movie, and the images were binarized to extract the pupil pixels. The pupil diameter was then determined as the maximum Feret diameter of the segmented pupil pixels for each frame. Frames with undetectable pupils due to blinking were excluded using a custom-written deviant detection program. Pupil sizes were normalized against the minimum and maximum diameters observed throughout the entire multi-day recordings for each mouse.

<u>Analysis of two-photon calcium imaging data</u>

Lateral motion was corrected using cross-correlation-based image alignment. Regions of interest (ROIs) corresponding to individual cell bodies or axon boutons were detected automatically using Suite2P software (*101*) and supplemented by manual drawing. All ROIs were individually inspected and edited for appropriate shapes using a custom graphical user interface in Matlab. Pixels within each ROI were averaged to derive a fluorescence time series, $F_{cell\text{-}meausred}(t)$. To correct for background contamination, ring-shaped background ROIs (starting at 2 pixels and ending at 8 pixels from the border of the ROI) were created around each cell ROI. From this background ROI, pixels that contained cell bodies or processes from neighboring cells (detected as pixels showing large increases in dF/F uncorrelated to that of the cell ROI during the entire imaging session) were excluded. The remaining pixels were averaged to generate a background fluorescence time series $F_{background}(t)$. The fluorescence signal for each cell was estimated as $F(t) = F_{cell\_measured}(t) - 0.9 \times F_{background}(t)$. To ensure robust neuropil subtraction, only cell ROIs that were at least 3% brighter than their corresponding background ROIs were included. To compensate for slow drifts in baseline fluorescence intensity, a normalized trace, $F_{norm}(t)$, was calculated as $F(t)/F_0(t)$, where $F_0(t)$ represents a time-varying drifting trace estimated by smoothing the inactive portions of F(t) using an iterative procedure.

The total receptive fields of individual cells were measured using their responses to 1-second pure tones. Cells were judged as significantly excited if they fulfilled two criteria: (1) $F_{norm}(t)$ had to exceed a fixed threshold value consecutively for at least 0.5 seconds in more than half of the trials, and (2) $F_{norm}(t)$ averaged across trials had to exceed the same fixed threshold value consecutively for at least 0.5 seconds. The threshold for excitation (3.0×SD during the baseline period) was determined using receiver operating characteristic (ROC) analysis to yield a 90% true positive rate in tone responses, while the threshold for inhibition (−1.5×SD) was set at half of that for excitation. Response amplitudes were quantified using a 1.2-second window after the tone onset. Best frequencies (BF) were identified as the frequency evoking the strongest excitatory response, independent of tone intensity. Bandwidth (BW70) was calculated as the average of the range of frequencies that evoked significant responses and the range of frequencies with a Gaussian fit exceeding a threshold at 70 dB SPL.

Cellular activity for multi-day experiments was assessed for significant excitation or inhibition based on two criteria: (1) the area above baseline for individual trials was significantly larger (or smaller for inhibition) than zero, using a two-sided Wilcoxon signed-rank test, and (2) the peak amplitude of the response exceeded 1×SD for excitation (or –0.5×SD for inhibition). For axon bouton imaging, a peak amplitude threshold of 0.2×SD was used for excitation, and inhibition was not assessed due to the low baseline fluorescence values. Given variations in tone durations (5, 7, and 9 seconds), statistical analyses focused on the first 5 seconds of tone presentation. Trial-averaged sound response traces for each cell were compiled from responses across the three tone durations. The area above (or below for inhibition) the baseline of the average trace during the first 5 seconds was used to calculate the Change Index (CI) as $CI = (Area_{after} - Area_{before}) / (Area_{after} + Area_{before})$. CIs were calculated separately for excitatory and inhibitory responses, and only responsive cells were included in the analysis. For example, if the CI was calculated from Day 1 to Day X, only cells that were responsive on Day 1, Day X, or both were included. For across-day CI time course plots, responses on each day were compared to those on Day 1. In muscimol infusion experiments, responses on Day 6 were compared to those on Day 5. In single-day experiments, averaged traces were calculated for blocks of 25 trials, and responses were compared to those in Block 1.

High-dimensional analysis of calcium imaging data

To compare within-day and across-day changes in the response patterns of A1 neuronal ensembles, we analyzed A1 population activity in a high-dimensional space, with each dimension corresponding to the response magnitude of individual neurons. We included only ROIs that were consistently visible across all imaging sessions and showed significant excitatory or inhibitory responses to tones in at least one session. Responses in each ROI were normalized to their minimum and maximum values across all sessions, resulting in normalized values that ranged from 0 to 1. We concatenated time series data from all trials across sessions to create an ensemble response matrix structured as ROIs × frames × trials. Although tone durations varied between 5, 7, and 9 seconds, only the first five seconds of tone presentations, shared across all trials, were included in the analysis. Direction vectors, representing plasticity in ensemble response patterns, were calculated for within-day ($Mode_{within}$) and across-day ($Mode_{across}$) changes. To ensure balanced contributions from onset-locked and sustained activities, we calculated a neuron's response magnitude in a trial by summing its normalized activity from 0.5–1.5 seconds and 4–5 seconds after sound onset. $Mode_{within}$ was obtained by computing the vector from the mean of Trials 1–10 to the mean of Trials 91–100 for each day and averaging these vectors over Days 1 to 5. $Mode_{across}$ was derived by calculating trial-averaged responses for Days 1 and 5 and computing their differences. The entire ensemble response matrix was projected onto $\text{Mode}_{\text{within}}$ and $\text{Mode}_{\text{across}}$ to visualize the within-day and across-day habituation of population response patterns. The kinetics of within-day plasticity were visualized by computing the difference between the ensemble activity in Trial 10 and that in Trial 1, and projecting this difference onto the $\text{Mode}_{\text{within}}$ axis. Similarly, the kinetics of across-day plasticity were visualized by computing the difference between the trial-averaged ensemble activity on Day 5 and Day 1, and projecting this difference onto the $\text{Mode}_{\text{across}}$ axis. These kinetics were quantified using the Ramp-up Index, which is defined as the ratio of the plasticity trace at 1 second and that at 5 seconds after tone onset. For the 6-day muscimol experiments, data from Day 6 (post-muscimol infusion) were projected onto $\text{Mode}_{\text{within}}$ and $\text{Mode}_{\text{across}}$, which were calculated based on data from Days 1 to 5.

NMDA receptor knockout experiments

To selectively ablate NMDA receptors from the A1 region, AAV8-syn-mCherry-Cre was co-injected with AAV9-syn-GCaMP6s into the A1 of $\text{Grin1}^{\text{flox/flox}}$ mice during the window implantation surgery. Following the injections and window implantation, mice were allowed to recover in their home cages for at least five weeks to ensure a sufficient reduction in NMDA receptor expression before habituation experiments. During two-photon calcium imaging, neurons expressing Cre were identified by their mCherry fluorescence. Control groups included both $\text{Grin1}^{\text{flox/flox}}$ mice without Cre virus injections and $\text{Grin1}^{+/+}$ mice with Cre virus injections. For targeted NMDA receptor knockout in specific inhibitory neuron subtypes, SST-Cre or VIP-Cre mice were crossed with $\text{Grin1}^{\text{flox/flox}}$ mice. The procedures for virus injections and surgeries were identical to those used for two-photon imaging in C57BL/6J mice. Mice were at least eight weeks old at the start of habituation experiments to ensure sufficient reduction in NMDA receptor expression. Data from mice carrying both homozygous and heterozygous alleles of floxed *Grin1* were pooled, as no significant differences were found between these genotypes. Control mice included $\text{Grin1}^{\text{flox/flox}}$ mice without *Cre* transgenes, SST-Cre mice without floxed *Grin1* transgenes, and VIP-Cre mice without floxed *Grin1* transgenes.

Electrophysiology

For single-unit recordings in the frontal cortex, mice were first implanted with a custom stainless-steel head bar and a cement cap. For recordings in habituated mice, the animals were head-fixed and exposed to prolonged 10 kHz pure tones (70 dB SPL, 5, 7, or 9-second durations, 7–16-second intervals) for 200 trials per day over five days prior to the recording. On the recording day, following a small craniotomy (<0.3 mm) and durotomy in the right frontal cortex, mice were head-fixed in an awake state, and a 384-channel probe (Neuropixels 1.0, imec) was slowly inserted (approximately 2 μm per second) perpendicular to the brain surface. To simultaneously record from OFCvl and M2, the probe was stereotaxically targeted to the coordinates (A, L, V) = (2.6, 0.75, 2.8). Additional probe insertion sites sampled a range of stereotaxic locations to record from surrounding frontal regions, including ACC, FRP, PL, IL, and OFCm. A reference electrode was placed at the dura above the frontal cortex of the contralateral hemisphere or the somatosensory cortex of the ipsilateral hemisphere. The probe was allowed to settle for at least one hour before collecting data. Unit activity was amplified, digitized, and acquired at 30 kHz using the OpenEphys system (https://open-ephys.org). During recordings, the mice sat quietly in a loosely fitted plastic tube in a sound-attenuating enclosure (Gretch-Ken). The tube was lined with fleece fabric for comfort and attenuation of noise from scratching. In two of the habituated mice, recordings were repeated over two days to sample multiple brain regions.

An intersectional viral approach was used to selectively activate OFCvl neurons projecting to A1 L2/3. Following the initial transcranial mapping of auditory cortical areas using intrinsic signal imaging in C57BL/6J mice, two small craniotomies (<300 μm) were performed over the low- and high-frequency domains of the right A1, where AAVretro-syn-Cre was injected (200–250 μm deep, 40 nL/site at 10 nL/min). In the same surgery, the right OFCvl was stereotaxically targeted for the injection of AAV8-

nEF-$C_{on}$/$F_{off}$-ChRmine-oScarlet (300 nL/site at 20 nL/min) at two coordinates: (A, L, V) = (2.4, 0.75, 2.7) and (2.8, 0.75, 2.65). A stainless steel head bar was cemented to the skull, and the skull's top surface was covered with black cement, leaving the areas around the craniotomies. The craniotomies were sealed with Duragel (Cambridge Neurotech), covered with a thin layer of transparent cement, and protected by an additional layer of silicone. The mice were allowed to recover in their home cages for 2-3 weeks to achieve the appropriate level of ChRmine expression. On the day of recording, following a small craniotomy (< 0.3 mm) and durotomy in the 10 kHz-responding domain of the A1, mice were head-fixed in an awake state, and a 384-channel probe was slowly inserted (approximately 1 μm per second) perpendicular to the brain surface. Spikes were monitored during probe insertion, and the probe was advanced until it reached the white matter, where no spikes were detected. The reference electrode was placed at the dura above either the visual or somatosensory cortex. A fiber-coupled 638 nm laser (RLM638TA-200FC; Shanghai Laser) was positioned 1-2 mm above the transparent cement covering the frontal cortex. ChRmine-expressing neurons were activated by transcranial illumination (1-second duration, continuous pulse, 50 Hz, 10 ms ramped pulses, or 20 Hz sinusoidal pulses; maximum intensity of 80 mW at the fiber tip outside the skull) at a 10- or 20-second interval. Control experiments were conducted with mice that did not express ChRmine. Successful transcranial activation of OFCvl units was confirmed by simultaneous recording from OFCvl during optogenetic activation.

Analysis of unit recording data

Single- and multi-units were isolated using Kilosort 2.5 software (*102*) and the spike-sorting graphical user interface Phy (*103*). Single-unit isolation was confirmed based on the consistency of the spike waveform and the inter-spike interval histogram. Specifically, after correction for the overall spike frequency, less than 5% of the spikes should fall within the refractory period, which was 2 ms and 1.5 ms for regular-spiking and fast-spiking units, respectively. The initial automated classification was conducted using the Bombcell spike classification toolbox (*104*). Following this automated step, spike clusters were individually inspected and edited to eliminate artifacts and duplications. Multi-unit spikes were calculated by aggregating all spikes within each channel or layer bin.

Post hoc identification of probe trajectories in the frontal cortex was achieved by examining DiI or DiO fluorescence in brain sections counterstained with DAPI. Brain sections were aligned to the Allen Common Coordinate Framework using Matlab codes modified from an existing repository (https://github.com/petersaj/AP_histology). Channels in the ventrolateral and lateral regions of OFC were collectively referred to as OFCvl. For A1 recordings, the positions of the cortical surface, layer borders, and white matter were identified using current source density analysis, the distribution of multi-unit spikes, and the correlation coefficient across channels (*105*).

Single units were considered significantly excited by tones if their spike rates during tone presentations were significantly higher than those during baseline period (5 seconds before the tone onset), as determined across trials using a two-sided Wilcoxon signed-rank test. Out of the 200 trials, the first 10 trials were excluded from analysis to remove the effect of rapid within-day habituation. In each mouse, the fraction of excited single units was calculated only for areas containing at least 30 isolated single units.

Peristimulus time histograms (PSTHs) were constructed using a 500-ms bin width. For visualization in Fig. 4K and fig. S8, PSTHs were smoothed using a Gaussian kernel. Tone response magnitudes in individual units were quantified by calculating the total spikes during tone presentations and subtracting the baseline firing rate obtained from the averaged PSTH. The effects of optogenetic manipulation were quantified using the Modulation Index (MI), which was calculated as (L-B)/(L+B), where L represents the activity during laser stimulation, and B represents the baseline activity. The MI values range from -1 to 1, where -1 indicates a complete loss of activity, 1 indicates the emergence of activity from nothing, and 0 indicates no change.

Anatomical tracing

Cell type-nonspecific retrograde tracing from A1 was conducted using two approaches: injecting AAVretro-syn-Cre in Ai9 mice or AAVretro-CAG-EGFP in VGAT-Cre × Ai9 mice. Following transcranial mapping with intrinsic signal imaging, the viruses were injected into the low-to-mid-frequency domain of A1 using a sharpened glass pipette (250 μm or 750 μm deep, 20 nL at 10 nL/min). Data from injections at both depths were combined, as no clear difference was observed in the distribution of presynaptic neurons.

For trans-synaptic retrograde tracing from specific neuron subtypes in A1, we used rabies virus-mediated monosynaptic tracing (*106*). Following transcranial mapping, a mixture of AAV5-CAGGS-Flex-Rev-mKate-T2A-CVS-N2cG and AAV5-CAG-Flex-Rev-mKate-T2A-TVA (for Emx1-Cre, SST-Cre, and PV-Cre-transgenic mice), or AAV9-CAGGS-Flex-Rev-mKate-T2A-CVS-N2cG and AAV9-CAG-Flex-Rev-mKate-T2A-TVA (for VIP-Cre mice) was injected into the mid-frequency domain of A1 (250 μm deep, 40 nL, 10 nL/min). Two weeks later, EnvA-pRbv-CVS-N2c(ΔG)-H2B-EGFP was injected at the same location (400 nL, 20 nL/min), resulting in the visualization of mKate$^+$/EGFP$^+$ starter cells around the injection site and EGFP$^+$ presynaptic cells in the frontal cortical areas.

Cell type-nonspecific tracing from the OFCvl was conducted by injecting AAV5-syn-EYFP in C57BL/6J mice for anterograde tracing and by injecting AAVretro-syn-mCherry in C57BL/6J mice or AAVretro-syn-Cre in Ai9 mice for retrograde tracing. The viruses were stereotaxically injected at the coordinates (A, L, V) = (2.7, 0.85, 2.7) (60–80 nL at 10 nL/min).

Two weeks (AAV) or one week (rabies) after virus injection, cardiac perfusion was performed using PBS, followed by 4% paraformaldehyde in PBS under isoflurane anesthesia. The brains were then extracted, fixed overnight with 4% paraformaldehyde, and immersed in 30% sucrose in PBS overnight for cryoprotection. The brains were coronally sectioned at 40 μm thickness using a freezing microtome

(SM2010R, Leica). After the sections were mounted on slide glasses and counterstained with DAPI, images were acquired using a fluorescence microscope (Axio Observer 7, ZEISS). Brain section images were aligned to the Allen Common Coordinate Framework using ABBA software (https://abba-documentation.readthedocs.io/en/latest/) to delineate anatomical area borders. The somas of retrogradely labeled presynaptic neurons were automatically detected and sorted into brain regions using QuPath software (https://qupath.github.io/). Histology figure panels were created by overlaying signals from multiple colors using Fiji (https://imagej.net/Fiji) or ZEN Microscopy (ZEISS) software.

Immunohistochemistry

For NMDA receptor immunostaining, brain slices were incubated in PBS containing 0.1% Triton X-100 and 10% normal donkey serum on a shaker for 1 hour at room temperature. The slices were rinsed three times with PBS for 5 minutes each and incubated in PBS with rabbit anti-NR1 primary antibodies (1:100, ab52177, RRID: AB_2112161, Abcam) on a shaker overnight at 4°C. The slices were then rinsed three times in PBS and incubated in PBS with Alexa 488-conjugated donkey anti-rabbit secondary antibodies (1:200, 711-545-152, RRID: AB_2313584, Jackson ImmunoResearch) for 2 hours at room temperature. After rinsing three times in PBS, the slices were mounted on slide glasses.

Statistics

Sample sizes were not predetermined by statistical methods but were selected based on standards commonly used in the field. All key results were replicated across multiple mice. Mice were randomly assigned to experimental groups. While the experiments were not performed blind, the experimenters were blind to the types and timings of trials during the ROI segmentation in two-photon imaging and spike sorting in electrophysiology. Data are presented as individual data points or mean ± SEM, as specified in the figure legends. Statistical differences between conditions were evaluated using standard two-sided non-parametric tests, specifically the Wilcoxon rank-sum test for unpaired data and the Wilcoxon signed-rank test for paired data. A Bonferroni correction was applied for multiple comparisons, and corrected p-values were reported. Fractions were compared using Fisher's exact test. Correlation coefficients were calculated as Spearman's correlation using Matlab's corr function. Two-way ANOVA was used to evaluate the effects of two independent variables.

**Supplementary Text:**

<u>Across-day habituation is not explained by global brain state changes</u>

We considered whether the weakening of sound responses with repeated exposure reflects a non-specific decrease in A1 responsiveness or is specific to the experienced sound. To address this, we assessed the difference between responses to two sound frequencies (Tone A and Tone B, chosen to flank the population best frequency of the imaged area). Tone A was presented daily, while Tone B was only encountered on the first and last days (fig. S3A). On Day 6 (post-habituation), A1 pyramidal neurons showed significantly decreased excitatory responses and increased inhibitory responses to Tone A (fig. 3, B to E). In contrast, Tone B responses on Day 6 remained at amplitudes equivalent to those before Tone A exposure, confirming the sound specificity of A1 sensory habituation (Tone A vs. Tone B, Block 1 excitation: $P = 8.4 \times 10^{-14}$, suppression: $P = 2.1 \times 10^{-10}$, two-sided Wilcoxon rank-sum test. Note that within-day habituation brought Tone B responses lower compared to the pre-exposure level when all 100 trials were considered). Together with the pupillometry results, these data demonstrate that across-day sensory habituation in A1 is due to sound-specific plasticity, rather than a general reduction in neural responsiveness due to global brain state changes.

<u>Multiple mechanisms for sensory habituation across time scales</u>

Previous studies have reported multiple mechanisms for short-term adaptation along the ascending auditory pathways. At the periphery, synaptic depression between hair cells and auditory nerves (*107*, *108*) causes rapid adaptation of auditory nerves and cochlear nucleus neurons to repetitive click trains on a millisecond timescale (*109*, *110*). More centrally, stimulus-specific adaptation (SSA) on a timescale of hundreds of milliseconds to seconds has been extensively studied in the inferior colliculus (*19*, *111*, *112*), medial geniculate body (*18*, *112*, *113*), and auditory cortex (*20*, *21*). SSA in the auditory cortex is likely due to a combination of inheritance from subcortical systems, short-term depression at thalamocortical synapses (*14*), and the short-term dynamics of cortical inhibitory neurons (*15*, *114*).

Although within-day habituation in our study operates on a shorter timescale than the OFC-dependent across-day habituation, its timescale (inter-stimulus intervals of 7–16 seconds) is still longer than the typical SSA and is comparable to the "adaptation for the long-term stimulus metastatistics" reported previously (*21*). Whether mechanisms similar to SSA apply to this adaptation at a middle-range timescale (seconds to tens of seconds) remains a topic for future research. It is plausible that longer timescales and greater complexity in stimulus statistics engage higher brain areas and more global network mechanisms in sensory adaptation. The involvement of frontal top-down circuits in across-day habituation represents the mechanism with the longest timescale within this spectrum of diverse adaptation mechanisms.

### Roles of novelty and prediction-error signals in sensory processing

Although our data do not support the role of novelty and prediction error signals in across-day sensory habituation, we do not rule out their potential influence on sensory processing under different conditions. Experimental paradigms specifically designed to create discrepancies between expected and actual sensory inputs have revealed prediction error responses in multiple cortical areas (*48*, *115*, *116*). In the auditory system, the effects of prediction error signals have been investigated using an oddball paradigm, which involves presenting standard sounds repeatedly alongside rare deviant sounds. In this setting, neural responses to predictable standard sounds adapt over hundreds of milliseconds (SSA), while stronger responses to unpredictable deviant sounds are observed as mismatch negativity (*117*). The potential involvement of top-down regulation in these prediction error responses has been suggested across multiple stages of auditory processing, including pathways from the auditory cortex to the inferior colliculus (*118*), from the higher-order auditory cortex to the primary auditory cortex (*119*), and from the prefrontal cortex to the auditory cortex (*120*). Nevertheless, the precise causal role of frontal cortical areas in modulating the auditory cortex through prediction or prediction error signals remains unclear.

In this study, we demonstrated a critical role for predictive signals within the OFC in filtering A1 activity based on daily experiences, consistent with the OFC's function of tracking predictive relationships within the external environment (*1*, *2*). On the other hand, several studies have reported reward prediction error signals in the OFC (*68*, *121–123*) (but see (*124–126*)), which may influence sensory processing in conditions with more explicit discrepancies between expected and actual sensory inputs. Further research should investigate how prediction and prediction error signals from the OFC support learning across different cognitive functions and determine if specific subdomains or subpopulations within the OFC (*65*, *85*) are responsible for these processes.

Comparison with previous studies using optogenetic activation of OFC

In a previous study, optogenetic activation of top-down projections from the OFC in vitro evoked direct excitatory postsynaptic currents in both excitatory and inhibitory neurons in A1 (*49*). Although not significant, the excitatory currents were stronger in inhibitory neurons than in excitatory neurons, providing a potential synaptic basis for the suppression of A1 neurons. Interestingly, while the same study reported a current sink in A1 L2/3 with OFC activation in vivo and suggested its role in modulating A1 plasticity, it did not report the effects of OFC activation on A1 neuronal firing (*49*, *50*). In contrast, we found consistent suppression of A1 activity with the activation of the OFC→A1 pathway (Fig. 4, L to N). This difference may be due to the variations in the optogenetic stimulation paradigms used. The previous study used isolated brief (5 ms) pulses, whereas we used prolonged trains (1 second). This distinction is crucial as excitatory inputs onto SST neurons show strong facilitation with repeated activations, unlike the depressing excitatory synapses onto pyramidal or PV cells. Consequently, the impact of SST cell-mediated inhibition becomes stronger with prolonged inputs (*82*, *127*). Slowly ramping-up predictive signals conveyed by OFC top-down inputs (Fig. 4F) are ideally suited for recruiting SST cells and generating prolonged suppression in A1 circuits. Supporting this idea, continuous 0.5-second optogenetic activation of the OFC has been shown to activate SST but not PV or VIP cells in the primary visual cortex (*89*). Therefore, the OFC-dependent sensory habituation via SST cell activation may represent a shared mechanism across sensory modalities.

**Supplementary Figures:**

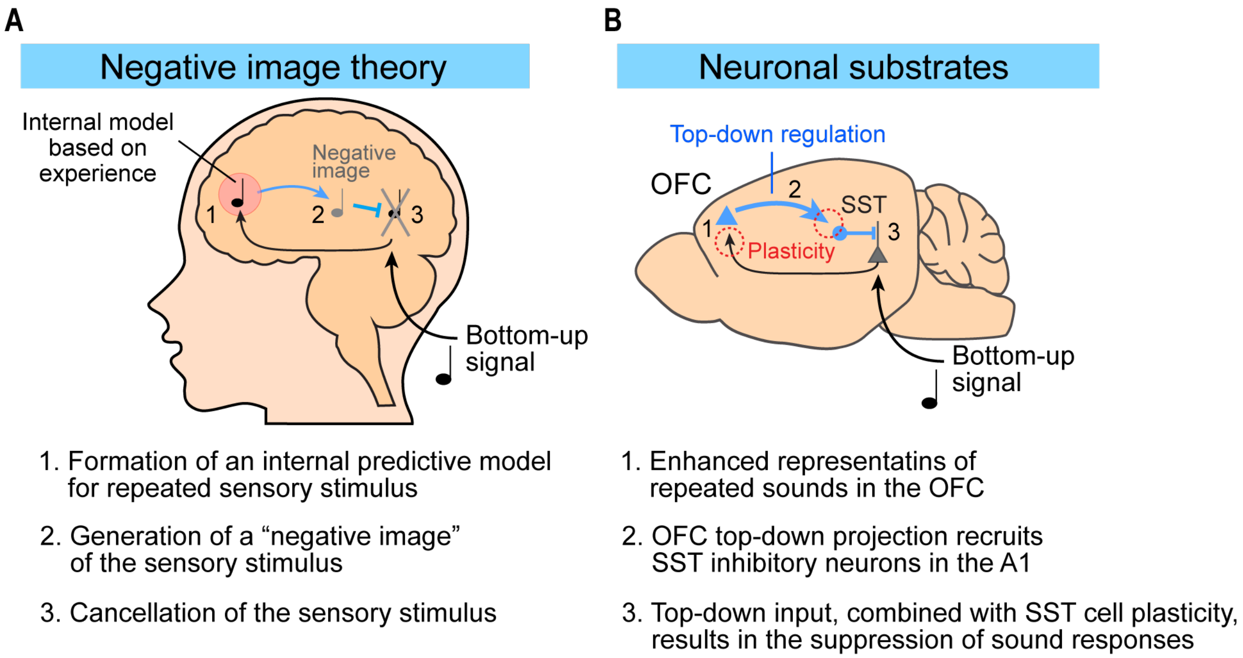


**Figure S1. Predictive negative image hypothesis for sensory habituation.** (**A**) Schematic illustrating the steps of sensory habituation mechanisms proposed by the predictive negative image hypothesis(*11*, *27*, *37*–*39*). The exact brain loci where the internal predictive model is stored and how this model filters sensory inputs remain unknown. (**B**) Summary diagrams of neuronal substrates for the negative image hypothesis proposed in this paper.

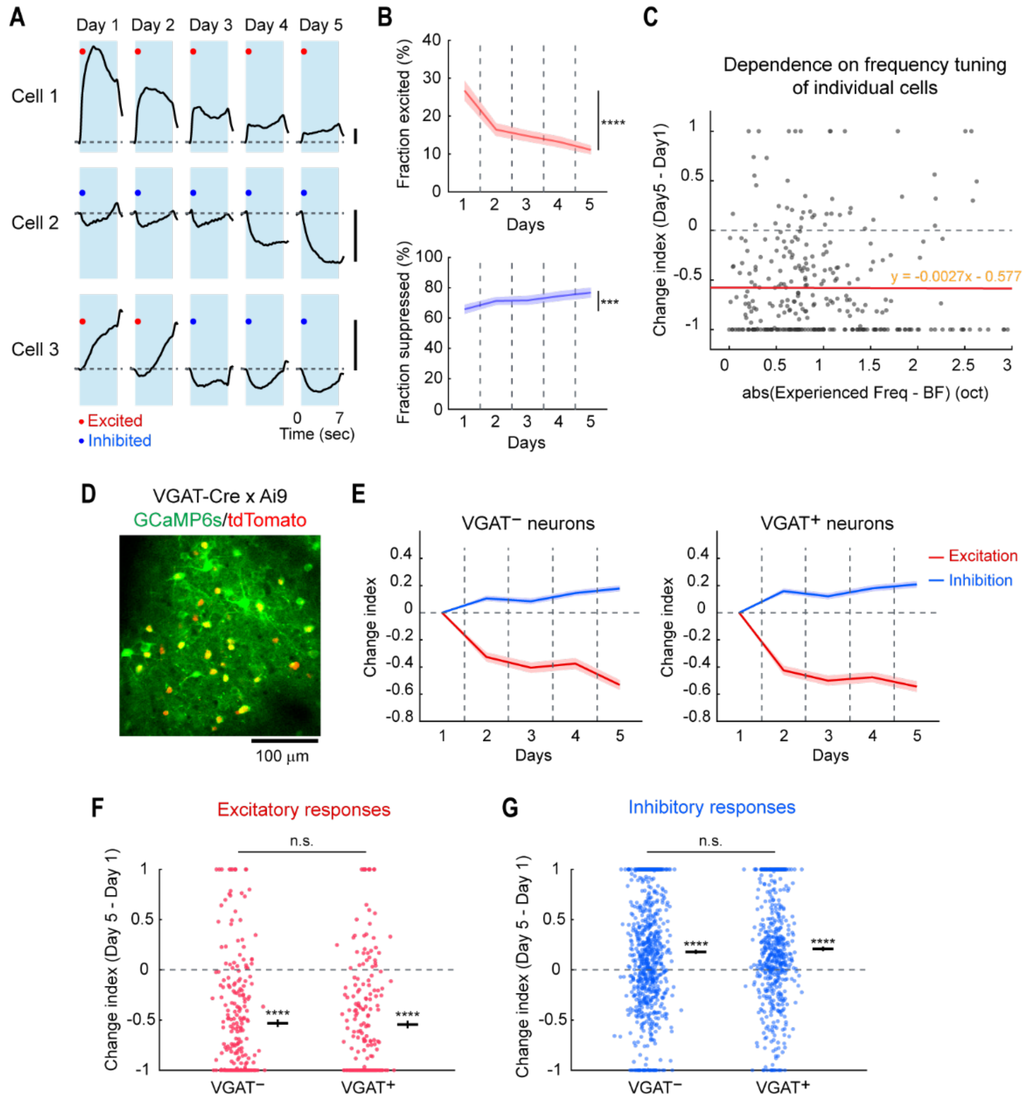


**Figure S2. Additional data characterizing A1 sensory habituation.** (**A**) Trial-averaged sound-evoked response traces from three cells tracked over five days. Red dots: significantly excited. Blue dots: significantly suppressed. Scale bars: 100% ΔF/F. (**B**) Fraction of significantly excited (top) and suppressed (bottom) cells over five days. *n* = 20 mice. ****$P = 8.9 \times 10^{-5}$, **$P = 2.5 \times 10^{-4}$ (two-sided Wilcoxon signed-rank test). Solid line: mean. Shading: SEM. (**C**) Change Index of excitatory responses from Day 1 to Day 5 in individual neurons plotted against the difference between their best frequency (BF) and the experienced frequency. Red line indicates a linear regression line. Sensory habituation occurs uniformly regardless of a cell's frequency tuning. *n* = 317 significantly responsive cells. $R = -0.0036$, $P = 0.95$ (Spearman correlation). (**D**) Representative two-photon image of A1 L2/3 neurons expressing GCaMP6s in all neurons and tdTomato in inhibitory neurons. (**E**) Change Index of excitatory (red) and inhibitory (blue) sound-evoked responses across days, plotted separately for $VGAT^-$ excitatory neurons (left) and $VGAT^+$ inhibitory neurons (right). *n* = 10 mice, 1,882 $VGAT^-$ neurons, 896 $VGAT^+$ neurons. (**F**) Change Index of excitatory responses from Day 1 to Day 5 in individual neurons, plotted separately for $VGAT^-$ and $VGAT^+$ neurons. $VGAT^-$, ****$P = 2.7 \times 10^{-64}$ (two-sided Wilcoxon signed-rank test), $VGAT^+$, ****$P = 7.8 \times 10^{-40}$; $VGAT^-$ vs. $VGAT^+$, $P = 0.51$ (two-sided Wilcoxon rank-sum test). (**G**) Same as (F) but for inhibitory responses: $VGAT^-$, ****$P = 4.7 \times 10^{-17}$; $VGAT^+$, ****$P = 8.5 \times 10^{-30}$; $VGAT^-$ vs. $VGAT^+$, $P = 0.27$.

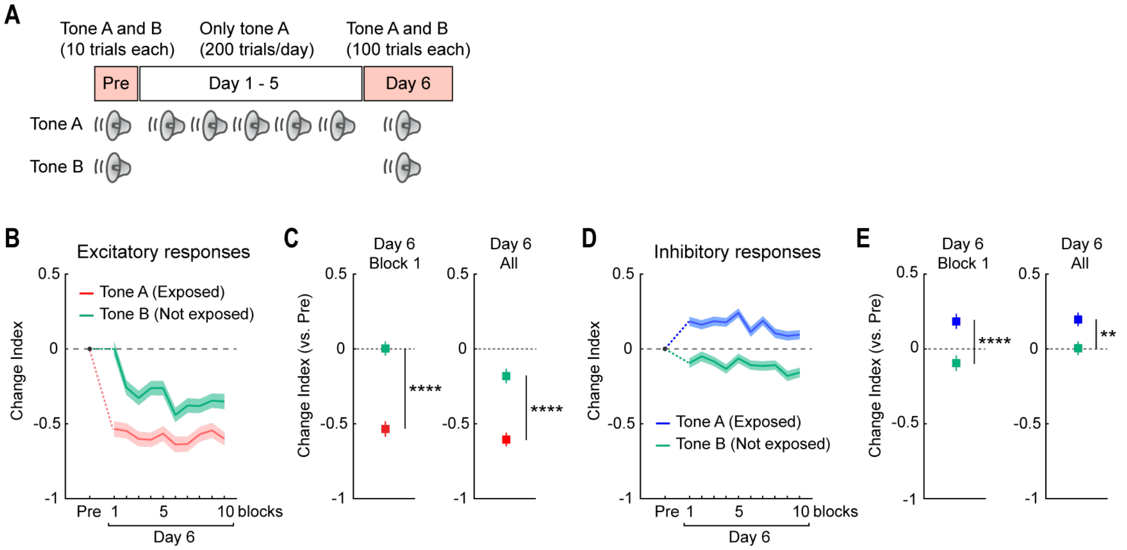


**Figure S3. Across-day habituation is selective to the exposed tone frequency.** (**A**) Schematic illustrating the sound-specific habituation paradigm using two sound frequencies. (**B**) Change Index of excitatory responses to Tone A (exposed) and Tone B (control) on Day 6 compared to the pre-habituation block. Each data point represents a block of 10 trials. $n$ = 1,040 cells from 4 mice. (**C**) Summary plots showing the Change Index of responses to Tones A and B in Block 1 of Day 6 (left) and all blocks of Day 6 (right). Tone A vs Tone B, Day 6 Block 1: ****$P = 8.4 \times 10^{-14}$ (two-sided Wilcoxon rank-sum test). Tone A vs. Tone B, All Day 6: ****$P = 3.6 \times 10^{-11}$. (**D-E**) Same as (B-C) but for inhibitory responses. Tone A vs Tone B, Day 6 Block 1: ****$P = 2.1 \times 10^{-10}$. Tone A vs. Tone B, All Day 6: **$P = 0.0084$.

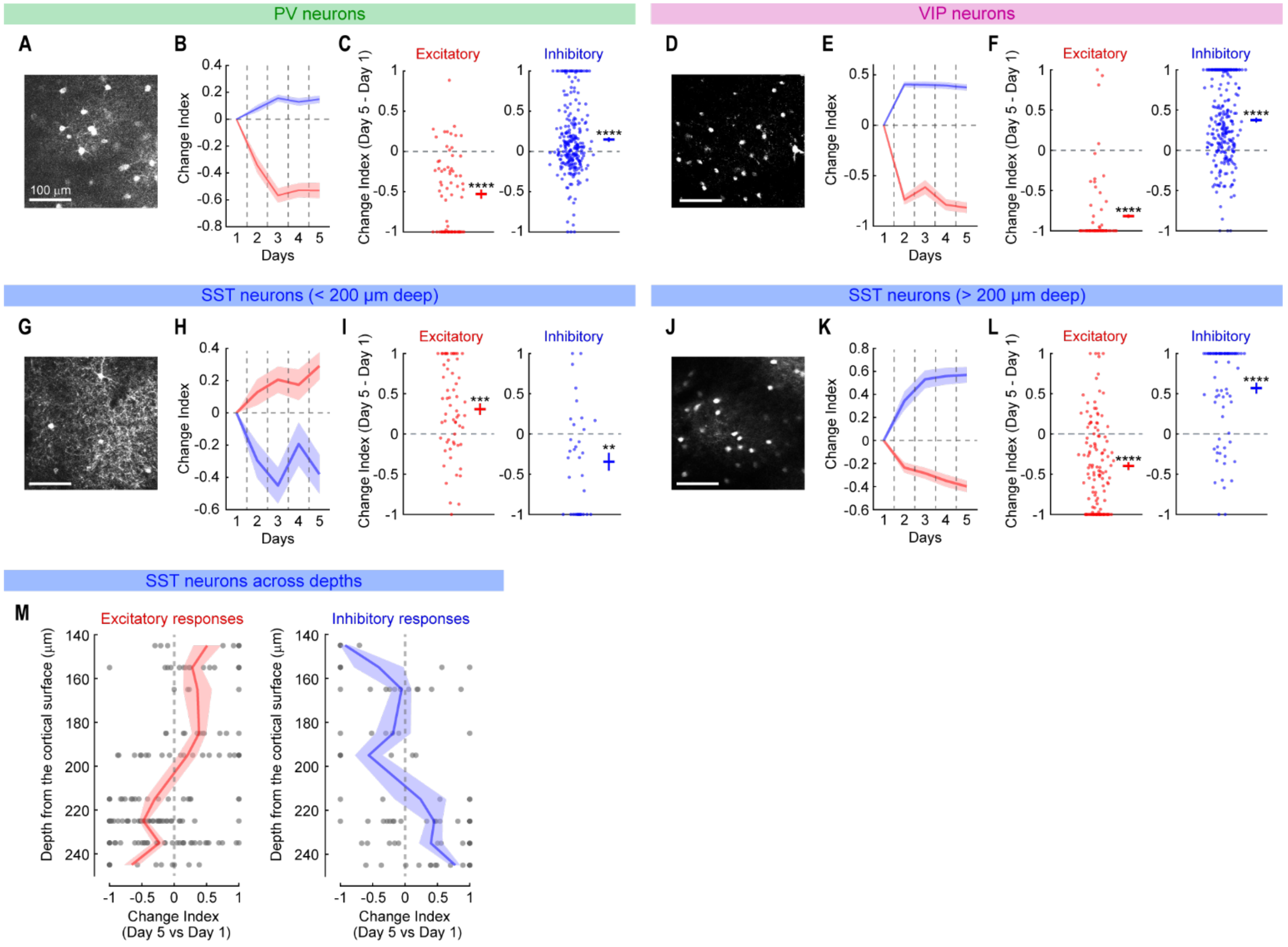


**Figure S4. Across-day habituation in three inhibitory neuron subtypes.** (**A**) Representative two-photon image of A1 L2/3 PV neurons expressing GCaMP6s. (**B**) Average (solid line) and SEM (shading) of the Change Index for excitatory (red) and inhibitory (blue) sound-evoked responses across days. *n* = 7 mice, 385 cells. (**C**) Change Index of excitatory (left) and inhibitory (right) responses in individual neurons on Day 5 compared to Day 1 across all mice. Lines on the right represent mean ± SEM. *n* = 71 significantly excited neurons and 265 significantly inhibited neurons. Excitatory, $****P = 2.2 \times 10^{-10}$ (two-sided Wilcoxon signed-rank test). Inhibitory, $****P = 2.0 \times 10^{-5}$. (**D-F**) Same as (A-C) but for VIP neurons. (**E**) *n* = 6 mice, 336 cells. (**F**) n = 70 significantly excited neurons and 268 significantly inhibited neurons. Excitatory: $****P = 1.1 \times 10^{-23}$. Inhibitory: $****P = 9.7 \times 10^{-26}$. (**G-I**) Same as (A-C) but for superficial (<200 μm from the surface) SST neurons. (**H**) *n* = 5 mice, 104 cells. (**I**) *n* = 60 significantly excited neurons and 34 significantly inhibited neurons. Excitatory, $***P = 6.3 \times 10^{-4}$. Inhibitory, $**P = 0.0022$. (**J-L**) Same as (A-C) but for deep (>200 μm from the surface) SST neurons. (**K**) *n* = 7 mice, 253 cells. (**L**) *n* = 126 significantly excited neurons and 76 significantly inhibited neurons. Excitatory, $****P = 3.8 \times 10^{-15}$. Inhibitory, $****P = 1.5 \times 10^{-6}$. (**M**) Laminar profile of the Change Index for excitatory (left) and inhibitory (right) sound-evoked responses in individual SST neurons on Day 5 compared to Day 1. Dots, individual cells. Solid lines and shading represent mean and SEM for data binned at 10 μm intervals. *n* = 12 mice.

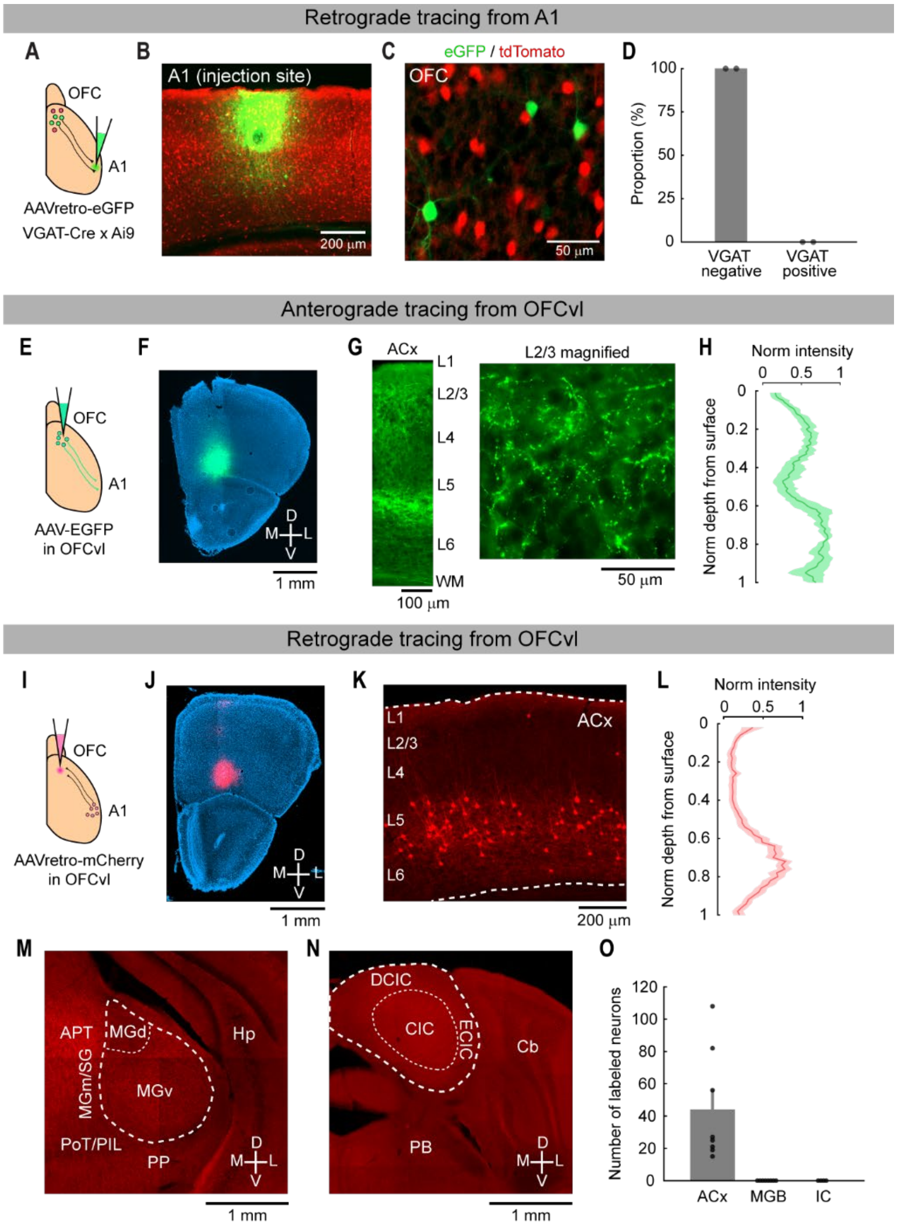


**Figure S5. Additional tracing data.** (**A**) Schematic illustrating two-color labeling of VGAT$^+$ inhibitory neurons (red) and retrogradely labeled neurons from A1 (green). (**B**) Coronal section around the injection site in the A1. (**C**) Magnified image of the OFCvl in a coronal section. There is no overlap between GFP$^+$ and tdTomato$^+$ neurons. (**D**) Summary bar plot quantifying the fraction of VGAT$^+$ and VGAT$^-$ cells out of retrogradely labeled neurons in the ipsilateral OFCvl. $n$ = 2 mice. (**E**) Schematic illustrating anterograde tracing from OFCvl. (**F**) Coronal section around the injection site in the OFCvl. (**G**) Left: Coronal section showing the laminar distribution of OFCvl axons in the auditory cortex (ACx). Right: Magnified image of L2/3. (**H**) Distribution of EGFP signal intensity across the normalized cortical depth. Solid lines and shadings show mean and SEM. $n$ = 4 sections from 2 mice. (**I**) Schematic illustrating retrograde tracing from OFCvl. (**J**) Coronal section around the injection site in the OFCvl. (**K**) Coronal section showing the OFCvl-projecting neurons in the auditory cortex. (**L**) Distribution of mCherry signal intensity across the normalized cortical depth. $n$ = 8 sections from 4 mice. (**M**) Coronal section showing no OFCvl-projecting neurons in the medial geniculate body (MGB). (**N**) Coronal section showing no OFCvl-projecting neurons in the inferior colliculus (IC). (**O**) Count of OFCvl-projecting neurons in the sections of the ipsilateral auditory cortex, MGB, and IC. *n* = 8 sections for ACx, 8 sections for MGB, 4 sections for IC, from 4 mice. APT, anterior pretectal nucleus; Cb, cerebellum; CIC/DCIC/ECIC, central nucleus/dorsal cortex/external cortex of IC; Hp, hippocampus; MGd/MGm/MGv, dorsal/medial/ventral subdivisions of MGB; PB, parabrachial nucleus; PIT, post intralaminar thalamic nucleus; PoT, post triangular thalamic nucleus; PP, peripeduncular nucleus; SG, suprageniculate thalamic nucleus.

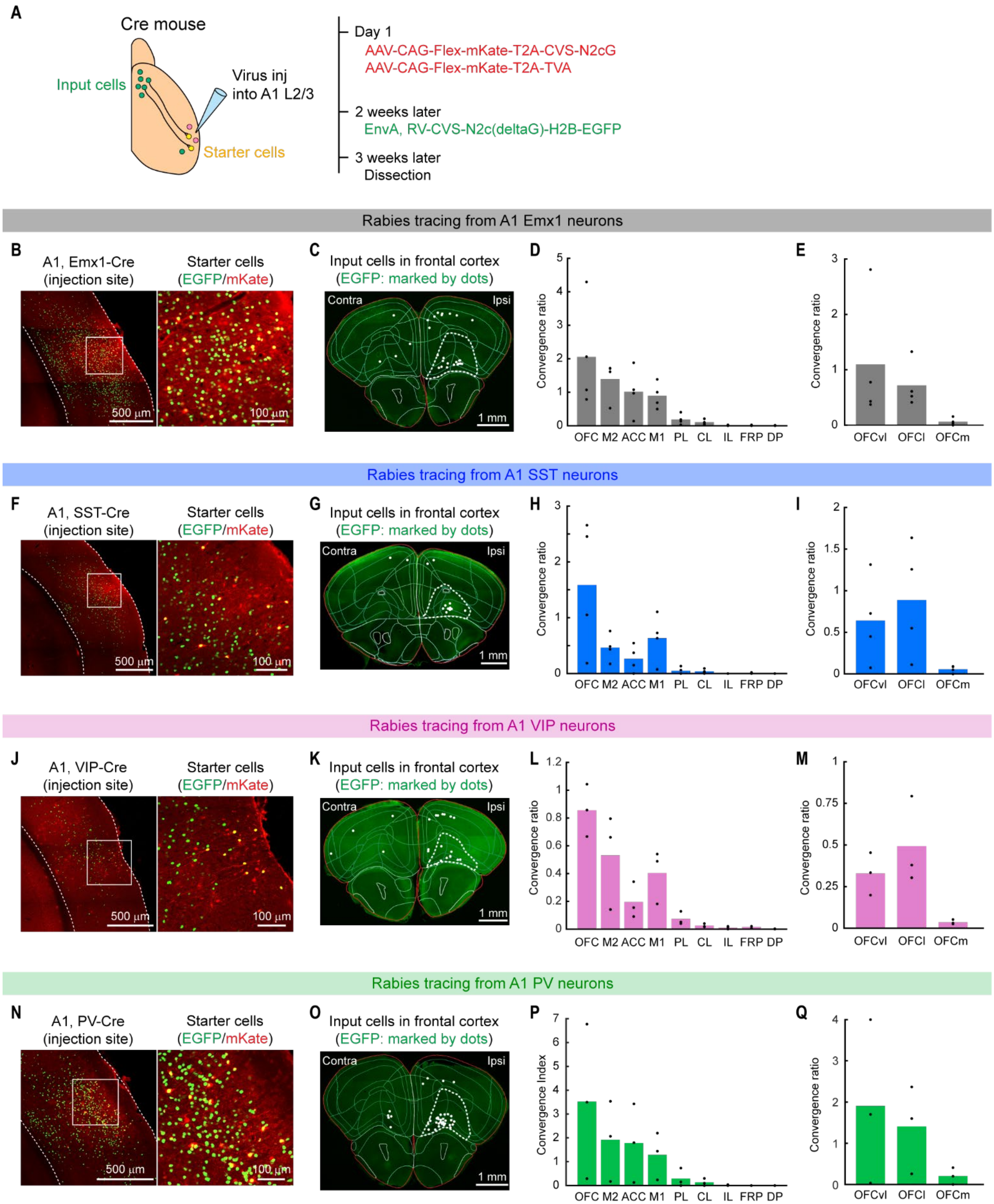

A
Cre mouse
Input cells
Virus inj
into A1 L2/3
Starter cells
Day 1
AAV-CAG-Flex-mKate-T2A-CVS-N2cG
AAV-CAG-Flex-mKate-T2A-TVA
2 weeks later
EnvA, RV-CVS-N2c(deltaG)-H2B-EGFP
3 weeks later
Dissection
Rabies tracing from A1 Emx1 neurons
B
A1, Emx1-Cre
(injection site)
Starter cells
(EGFP/mKate)
500 μm
100 μm
C
Input cells in frontal cortex
(EGFP: marked by dots)
Contra
Ipsi
1 mm
D
Convergence ratio
OFC M2 ACC M1 PL CL IL FRP DP
E
Convergence ratio
OFCvl OFCl OFCm
Rabies tracing from A1 SST neurons
F
A1, SST-Cre
(injection site)
Starter cells
(EGFP/mKate)
500 μm
100 μm
G
Input cells in frontal cortex
(EGFP: marked by dots)
Contra
Ipsi
1 mm
H
Convergence ratio
OFC M2 ACC M1 PL CL IL FRP DP
I
Convergence ratio
OFCvl OFCl OFCm
Rabies tracing from A1 VIP neurons
J
A1, VIP-Cre
(injection site)
Starter cells
(EGFP/mKate)
500 μm
100 μm
K
Input cells in frontal cortex
(EGFP: marked by dots)
Contra
Ipsi
1 mm
L
Convergence ratio
OFC M2 ACC M1 PL CL IL FRP DP
M
Convergence ratio
OFCvl OFCl OFCm
Rabies tracing from A1 PV neurons
N
A1, PV-Cre
(injection site)
Starter cells
(EGFP/mKate)
500 μm
100 μm
O
Input cells in frontal cortex
(EGFP: marked by dots)
Contra
Ipsi
1 mm
P
Convergence Index
OFC M2 ACC M1 PL CL IL FRP DP
Q
Convergence ratio
OFCvl OFCl OFCm

**Figure S6. Monosynaptic innervation of A1 SST cells by frontal cortical areas.** (**A**) Schematic illustrating the rabies viral strategies for monosynaptic retrograde tracing from Cre-expressing neurons in A1 L2/3. Red circles: mKate-expressing AAV-infected neurons. Green circles: EGFP-expressing rabies-infected neurons. Yellow circles: starter cells co-expressing mKate and EGFP. (**B-E**) Retrograde tracing from A1 L2/3 pyramidal neurons using Emx1-Cre mice. (**B**) Left: Coronal section of the injection site in A1. Dotted lines: cortical borders. Right: Magnified view of the area indicated by a square in the left image. (**C**) Coronal section of the frontal areas overlaid with area borders. White dots: presynaptic neurons. (**D**) Bar plots summarizing the distribution of input cells in ipsilateral frontal cortical areas. *n* = 4 mice. (**E**) Further classification of OFC into ventrolateral (OFCvl), lateral (OFCl), and medial (OFCm) subdivisions. (**F-I**) Same as (B-E) but from A1 SST neurons using SST-Cre mice (*n* = 4). (**J-M**) Same as (B-E) but from A1 VIP neurons using VIP-Cre mice (*n* = 3). (**N-Q**) Same as (B-E) but from A1 PV neurons using PV-Cre mice (*n* = 3). White dotted lines in the frontal cortical images indicate the boundary of the combined OFCvl and OFCl.

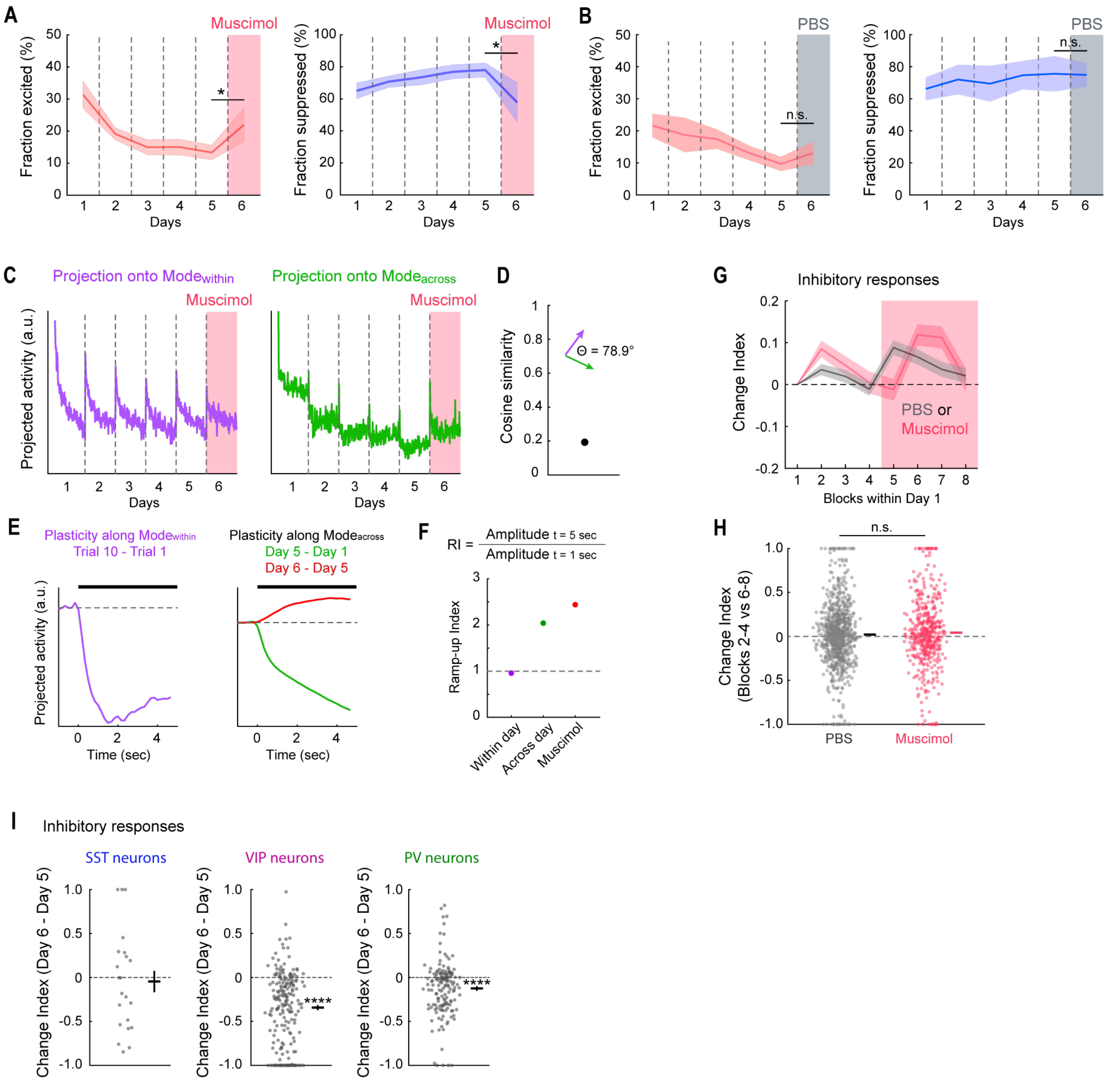


**Figure S7. Additional data characterizing the effect of OFC inactivation.** (**A-B**) Additional data from A1 L2/3 pyramidal cell imaging during muscimol infusion into the OFCvl. (**A**) Fraction of significantly excited (left) and suppressed (right) cells over six days. $n$ = 7 mice, 1,177 responsive cells. Excited: *$P$ = 0.031. Suppressed: *$P$ = 0.016. (**B**) Same as (a) but for PBS infusion into the OFCvl. $n$ = 3 mice, 439 responsive cells. Excited: $P$ = 0.25. Suppressed: $P$ = 1.0. (**C**) Projection of sound-evoked A1 ensemble trial-to-trial activity dynamics onto Mode$_{within}$ (left) and Mode$_{across}$ (right) defined using data from Days 1–5. Data points represent individual trials (100 trials × 6 days) before and after muscimol infusion. The ensemble activity pattern shows partial recovery along Mode$_{across}$ but not Mode$_{within}$ following muscimol infusion. $n$ = 7 mice, 734 cells imaged throughout the six days. (**D**) Cosine similarity between Mode$_{within}$ and Mode$_{across}$, indicating a nearly orthogonal relationship. (**E**) Left: Change in sound-evoked ensemble activity along Mode$_{within}$

between Trial 1 and Trial 10, illustrating prominent within-day, across-trial habituation around tone onset. Right: Change in sound-evoked ensemble activity along $\mathrm{Mode}_{\mathrm{across}}$ from Day 1 to Day 5 (green) and from Day 5 to Day 6 (red). Both across-day habituation and its recovery by muscimol slowly ramp up during the sustained tone. (**F**) Summary data showing the Ramp-up Index for across-day plasticity (Day 1 to Day 5), muscimol-induced recovery (Day 5 to Day 6), and within-day plasticity (Trial 1 to Trial 10). (**G**) Change Index for inhibitory responses across blocks of trials during muscimol infusion on Day 1 in naïve mice. Each data point represents a block of 25 trials. Red: muscimol ($n$ = 4 mice). Dark gray: PBS control ($n$ = 6 mice). (**H**) Change Index for inhibitory responses in individual neurons in blocks 6–8 compared to blocks 2–4 across all mice. $n$ = 894 and 492 significantly suppressed neurons for PBS and muscimol. $P$ = 0.55 (two-sided Wilcoxon rank-sum test). (**I**) Change Index for inhibitory responses in SST, VIP, and PV neurons comparing Day 5 and Day 6. SST, $n$ = 22 responses from 4 mice, $P$ = 0.30; VIP, $n$ = 197 responses from 5 mice, ****$P = 9.0 \times 10^{-28}$; PV, $n$ = 150 responses from 5 mice, ****$P = 2.7 \times 10^{-7}$ (two-sided Wilcoxon signed-rank test).

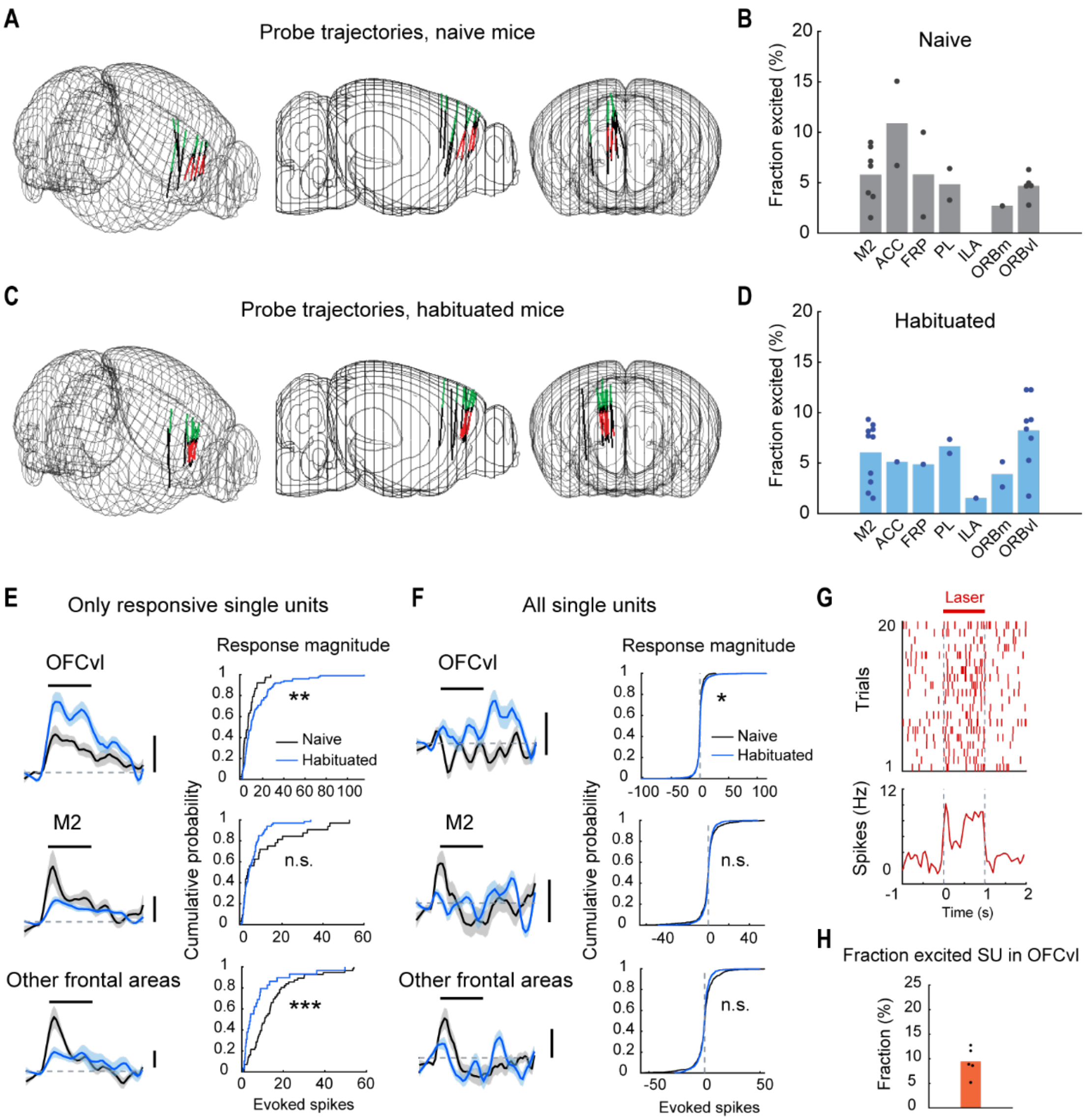


**Figure S8. Additional data characterizing sound-evoked responses in frontal areas.** (**A**) Summary of all probe penetrations in naïve mice aligned to the Allen Common Coordinate Framework. Probe tracks within M2 and OFC are shown in green and red, respectively. *n* = 11 penetrations from 8 mice. View angles from top-right, right, and front are shown. (**B**) Fraction of single units with significant excitatory responses in the frontal areas of naïve mice. Individual dots represent mice, and bar plot represents mean. (**C-D**) Same as (A-B) but for recordings in habituated mice. *n* = 9 penetrations from 9 mice. (**E**) Left: Sound-evoked response traces averaged across significantly excited single units in naïve (black) and habituated (blue) mice. Black bar represents the sound timing. Scale bar: 0.5 Hz. Right: Cumulative plots of sound-evoked spike counts in significantly excited single units. Top: OFCvl (*n* = 38 and 99 units for naïve and habituated; $^{**}P = 0.010$). Middle: M2 (*n* = 32 and 66 for naïve and habituated; $P = 0.55$). Bottom: other frontal areas, including ACC, FRP, PL, IL, and OFCm (*n* = 56 and 29 for naïve and habituated; $P = 9.8 \times 10^{-4}$). Two-sided Wilcoxon rank-sum test. OFCvl and M2 data are the same as Fig. 4K. (**F**) Same as (E) but for all single units, including non-responsive ones. Top: OCvl (*n* = 785 and 1203 units for naïve and habituated. $^{*}P = 0.027$. Middle: M2 (*n* = 632 and 1,059 for naïve and habituated; $P = 0.49$). Bottom: other frontal areas (*n* = 731 and 549 for naïve and habituated; $P = 0.054$). (**G**) Raster (top) and peristimulus time histogram (bottom)

of a representative single unit in OFCvl activated by a laser. (**H**) The fraction of OFCvl single units significantly excited by a laser (*n* = 5 mice).

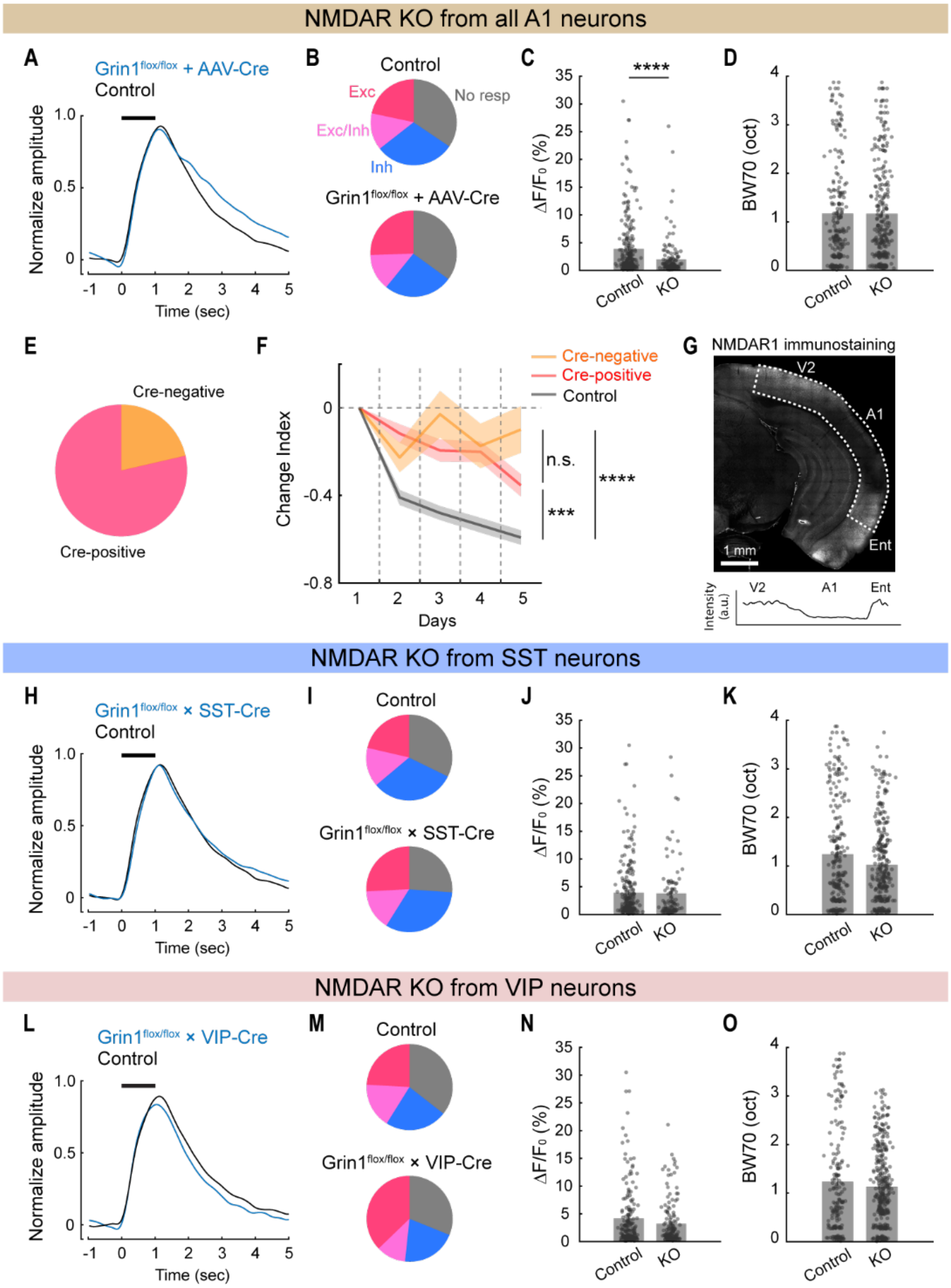


**Figure S9. Additional data characterizing NMDA receptor knockout experiments.** (**A-D**) Characterization of tonal receptive fields using one-second tone pips in mice with NMDA receptor knockout from A1. (**A**) Normalized response traces to the best frequency tone pips, averaged across all responsive neurons from Grin1$^{flox/flox}$ + AAV-Cre (KO; blue) and control (black) experiments. Bar, tone. (**B**) Fraction of cells with significant excitatory (Exc), inhibitory (Inh), and both excitatory and inhibitory (Exc/Inh) responses across mice. No resp: nonresponding cells. Control: *n* = 5 mice, 1,101 cells from 6 FOVs; KO: *n* = 4 mice, 1,275 cells from 5 FOVs. (**C**) Scatter plot showing the response magnitudes of neurons at their best frequencies. *n* = 236 and 143 responsive cells for control and KO mice. ****$P = 1.7 \times 10^{-6}$. Two-sided Wilcoxon rank-sum test. (**D**) Scatter plot showing the bandwidth of neurons at 70 dB SPL (BW70). *P* = 0.84. (**E**) Fraction of Cre-positive and Cre-negative neurons in the imaged fields of view, determined by their mCherry expression. *n* = 905 imaged neurons. (**F**) Grin1$^{flox/flox}$ + AAV-Cre data from Fig. 5B, categorized by the presence (red) or absence (orange) of Cre expression in imaged neurons. Reduced habituation is also observed in Cre-negative neurons, suggesting a network-level effect of habituation. *n* = 226, 154, and 42 for Control, Cre-positive, and Cre-negative. Control vs. Cre-positive: ***$P$ = 8.4 × $10^{-4}$. Control vs. Cre-negative: ****$P$ = 3.0 × $10^{-5}$. Cre-positive vs. Cre-negative: *P* = 0.091. Two-sided Wilcoxon rank-sum test with Bonferroni correction. (**G**) Top: Coronal section around the AAV injection site, showing NMDA receptor immunostaining. Bottom: Fluorescence intensity quantified along the dotted rectangle in the top image. (**H-K**) Same as (A-D) but for mice with NMDA receptor knockout from SST neurons. (**I**) Control: *n* = 6 mice, 243 cells from 6 FOVs. KO: *n* = 5 mice, 107 cells from 5 FOVs. **(J)** *n* = 243 and 107 responsive cells for control and KO mice. *P* = 0.56. (**K**) *P* = 0.14. (**L-O**) Same as (A-D) but for mice with NMDA receptor knockout from VIP neurons. (**M**) Control: *n* = 5 mice, 921 cells from 5 FOVs. KO: *n* = 4 mice, 1,256 cells from 6 FOVs. (**N**) *n* = 191 and 154 responsive cells for control and KO mice. *P* = 0.14. (**O**) *P* = 0.76.
